## Supplemental Information and Figures for "Hemibiotrophic fungal pathogen induces systemic susceptibility and systemic shifts in wheat metabolome and microbiome composition"

**Supplementary Materials and Methods**

**Fungal, bacterial, and plant material**

The Dutch *Zymoseptoria tritici* isolate IPO323 was kindly provided by Gert Kema (Wageningen University, The Netherlands). The fungus was maintained at 18°C either in liquid yeast-malt-sucrose (YMS) medium on a horizontal shaker or on solid YMS agar. *Pseudomonas syringae* pathovar *Por36_1rif* was kindly provided by Henk-Jan Schoonbeek (John Innes Centre, United Kingdom). *P. syringae* pathovars *Pst DC3000*, *Pst DC3000 hrcC-*, and *Pma ES4326* were kindly provided by Tiziana Guerra (Leibniz-Institut für Gemüse- und Zierpflanzenbau, Germany). All *P. syringae* strains were maintained at 28°C either in liquid Kings B medium (KB) on a horizontal shaker or on solid KB agar containing 50 mg/L rifampicin (KBrif). The *Triticum aestivum* cultivar Obelisk was obtained from Wiersum Plantbreeding BV (Winschoten, the Netherlands). Wheat cultivar Chinese Spring was kindly provided by Bruce McDonald (ETH Zurich, Switzerland). Plants were grown in phytochambers on peat under constant conditions with a day/night cycle of 16 h of light (~200 µmol/m2/s) and 8 h of darkness in growth chambers at 20°C and 90% relative humidity.

**Fungal infections**

*Z. tritici* strain IPO323 was grown on YMS solid medium for 5 days at 18°C before inoculation of liquid YMS medium and cell growth for another 3 days at 18°C. Fungal inoculum was adjusted to 1 x 107 cells/mL in $H_2O$ with 0.1% (v/v) Tween 20. Fungal inoculum, control treatment ($H_2O$ + 0.1% (v/v) Tween 20) or heat-killed fungal inoculum (1 x 107 cells/mL in $H_2O$ with 0.1% (v/v) Tween 20, incubated for 10 min at 96°C) was brushed onto labelled areas (6 cm) of the second leaf of each plant. Inoculated plants were placed in sealed bags containing water for 48 h to facilitate 100% relative humidity. Bags were removed after 48 h, and plants were kept in phytochambers until the end of the experiment. Upon completion of the experiment after 21 days, the necrotic leaf area and pycnidia formation was determined. Infected leaves were scanned at a resolution of 2400 dpi using a flatbed scanner (HP Photosmart C4580; HP, Böblingen, Germany). Scanned images were analyzed using automated image analysis and ImageJ1.

**Bacterial infections**

Analysis of *in planta* growth of *P. syringae* in wheat was adapted from Schoonbeek *et al.*[2]. *P. syringae* strains were grown for 24 h at 28°C on KBrif agar plates. Bacteria were resuspended in 5% KB liquid medium + 0.1% (v/v) Tween 20 to an $OD_{600}$ nm of 0.02. For wheat inoculation, four 1-mm holes (distance 1 cm each) were punctured in the labeled areas (6 cm) of the second leaf of 15-day-old wheat plants with a pin (if not stated otherwise). A 2-µL droplet of bacteria suspension or control treatment (5% KB + 0.1% (v/v) Tween 20) was applied on each hole, and droplets were allowed to dry. The plants were sealed in a plastic bag and incubated for 4 days in phytochambers. At 4 days post infection (dpi), disease symptoms appeared as yellow circles and were assessed by counting bacteria after extraction and serial dilution as follows. Leaf discs (cork borer size 1) of the area around the punctured holes of inoculated leaf were isolated and disrupted with 3 ceramic balls (2.8 mm) in a Precellys Tissue Homogenizer (4500 rpm, 10 sec) (Bertin Technologies, Frankfurt am Main, Germany) to release bacteria from the leaf apoplast. A 1:10 to 1:1000000 dilution series was made in 5% KB, and 10 µL from each dilution were spotted on KBrifcyc (KBrif + 50 mg/L cycloheximide) and incubated at 28°C for 24 to 36 h. The number of colonies was counted to calculate the number of colony forming units (CFUs) per $cm^2$.

For local bacterial coinfection experiments, the area for bacterial infection was the same area that was previously infected with IPO323. For adjacent bacterial coinfection experiments, the labeled area for bacterial infection was separated from the fungal infection area by a 1-cm buffer zone (Fig. S14). Systemic bacterial coinfection took place on the third leaf, while fungal spores were applied on the second leaf. If not stated otherwise, fungal infection took place 4 days before bacterial infection according to the aforementioned protocol.

**Systemic fungal coinfection and biomass assay**

We carried out the fungal infections in the systemic fungal coinfection experiment according to the aforementioned protocol. Local infection with *Z. tritici* strain IPO323 on the second leaf was followed by coinfection with the same strain on the third leaf 4 days after the initial infection of the second leaf. The second and third leaves were harvested at the indicated time points. Pictures for a visual overview of symptom development were taken (Sony Alpha 6000), and all leaves were scanned individually at a resolution of 2400 dpi using a flatbed scanner (HP Photosmart C4580; HP, Böblingen, Germany). Scanned images were analyzed for necrosis and pycnidia formation using automated image analysis and ImageJ[1]. After scanning, third leaves were immediately dipped into liquid latex. The latex was dried and then peeled from the leaves to remove extraneous fungal material from the leaf surface. The resulting leaf samples contained fungal cells only from inside the leaf tissue. Samples were deep-frozen, disrupted with 3 ceramic balls (2.8 mm) at -20°C in a Precellys Tissue Homogenizer (5500 rpm, 30 sec, 2 cycles, 15-sec break) (Bertin Technology SAS). After homogenization, the samples were stored at -80°C until further use. DNA was extracted using a standard phenol-chloroform extraction protocol[3]. Total DNA was quantified using a Qubit Fluorometer (Thermo Fisher Scientific, Dreieich, Germany). Fungal DNA was quantified by quantitative real time PCR (qPCR) (CFX96 System, Bio-Rad Laboratories, München, Germany) of fungal GAPDH with iQ SYBR Green (Bio-Rad Laboratories). A DNA standard curve was prepared from *Z. tritici in vitro* culture.

**Staining of hydrogen peroxide**

To detect $H_2O_2$, DAB (3,3′-diaminobenzidine) staining was carried out following an adaption of previously published protocols[4–6]. Infected leaf areas were collected at the indicated time points and infiltrated under gentle vacuum with 1 mg/mL DAB + 0.1% Tween 20 and 10 mM sodium phosphate buffer (pH 7). The staining (in darkness) was terminated after 90 min. Leaves were transferred into 75% (v/v) ethanol and 25% (v/v) acetic acid, fixed at room temperature until bleached completely, and immersed in 40% glycerol before further analysis.

**Metabolomics profiling**

**Metabolite extraction**

Leaf material (6-cm leaf area) was harvested, weighed, deep-frozen, and disrupted with 3 ceramic balls (2.8 mm) in a Precellys Tissue Homogenizer (5500 rpm, 30 sec, 2 cycles, 15-sec break) (Bertin Technologies, Frankfurt am Main, Germany). The homogenizer was chilled to -20°C to protect the metabolites for oxidation or temperature degeneration. After homogenization, the samples were stored at -80°C. When ready for use, 750 µL methanol/water solution (80/20, v/v; ultra LC-MS Rotisolv grade; Carl Roth, Karlsruhe, Germany) were added to each sample. The mixture was vortexed for 20 seconds and stored on crushed ice. Then, samples were placed in an ultrasonic bath (4°C) and lysed for 1 minute. After centrifugation (13000 x *g*, 5 min), the supernatant was transferred to a new tube. The pellet was re-extracted with 750 µL methanol/water solution, vortexed, and placed in an end-over-end shaker (1 h at room temperature). After centrifugation, the supernatant was transferred and combined with the first supernatant, mixed, and stored by -80°C until further analysis. For analysis, the samples were diluted 1:10 with a mixture of water/methanol/acetic acid (49.95/49.95/0.1 (v/v/v); ultra LC-MS Rotisolv grade; Carl Roth, Karlsruhe, Germany) and transferred to 1.5-mL HPLC vials.

**Metabolite measurements**

A Fourier-transform ion cyclotron resonance mass spectrometer (FT-ICR-MS; 7 Tesla, SolariXR, Bruker, Bremen, Germany) was used and was linked with an autosampler and pump from an HPLC 1260 Infinity

(Agilent, Waldbronn, Germany). The samples were stored at 4°C in the autosampler and automatically injected to the MS with a flow of 0.015 µL/min. Water/methanol (50/50, v/v) with 0.1% acetic acid was used as eluent, and 60 µL were injected. The samples were measured with ESI in the positive and negative ionization mode. The mass range was from 65 to 950 Daltons. The main instrument parameters were: dry gas (nitrogen) temperature of 200°C with 4 L/min; nebulizer (nitrogen) 1 bar; time-of-flight time section 0.35 ms, and the quadrupole mass was 150 m/z with a RF frequency of 2 MHz; and detector sweep excitation power of 18%. The average resolution for the mass 400 m/z was 600,000.

The data evaluation was conducted with DataAnalysis 5.0 and MetaboScape 4.0 (both Bruker, Bremen, Germany). Data were mass recalibrated versus the internal sample matrix (mostly free amino acids, sugars, and organic acids) with a tolerated error below 0.5 ppm. The importing parameters were tolerated mass error 0.5 mDa and an intensity threshold of 1,000,000 counts. Data were separately evaluated for each adduct. For each bucket, the sum formula was calculated by means of SmartFormula (Bruker, Bremen, Germany) and based on the 7 gold rules of Kind and Fiehn[7]. In addition, a second annotation was conducted with customized databases by suspected targeted methods. The database was created based on 15 different plant-related pathways (from KEGG[8–10]), such as the general secondary plant metabolism or phenylpropanoid biosynthesis pathways (Table S5).

**Statistical analysis of metabolomics dataset**

All m/z values with assigned sum formula were exported and further processed with the statistical software R (version 3.4.2). For each method (SM/UM) and mode (positive/negative), measurements of m/z values assigned to the same sum formula were added, and metabolites with measurements in fewer than 3 samples were removed. To create a single metabolomics data set, metabolite data of the 4 modes were merged. For metabolites that were available in several modes, the one with the smallest number of missing values, or if ambiguous the largest median, was kept. Metabolite measurements were $\log_2$ transformed, and missing values were replaced by the limit of detection (= $\log_2(10^6)$).

Metabolite measurements were compared separately between breeds, positions, and treatments, and each comparison was stratified by the 3 remaining factors (including day). A linear regression model was fit for each metabolite separately, and *P* values were corrected for multiple testing using the Benjamini-Hochberg procedure[11].

**Microbiota profiling**

**Plant growth for microbiota profiling**. Prior to sowing, seeds were washed 3 times (1 min each) with 20 mL sterile water, then 1% phosphate-buffered saline (v/v, PBS) supplemented with 0.02% (v/v) Tween 20, and then sterile 1% PBS. Two seeds per cultivar (i.e., *T. aestivum* cultivar Chinese Spring or

Obelisk) where sowed in a pot containing 155 g of peat (HAWITA GmbH, Vechta, Germany). The peat was preinoculated with ~70 mL of soil slurry that was obtained by diluting 1 g soil (Kiel agricultural soil collected from Hohenschulen experimental farm) in 100 mL sterile double distilled (dd) water. Pots were gathered in a tray, flooded with 2 L tap water, and covered with a plastic bag for 2 days. Plants were incubated in phytochambers at 20°C with 90% relative humidity with 16 hours of light for an additional 15 days prior to the fungal infection. Fungal infection was carried out according to the aforementioned protocol.

**Leaf sampling and DNA extraction.** Leaf samples were harvested at 0, 4, and 8 days after infection with *Z. tritici* (IPO323) in 50-mL sterile tubes separately for adjacent and local tissues. Each sample was washed 3 times as indicated above. Leaves were briefly dried with sterile blotting paper (0.35 mm, Hahnemühle, Germany), transferred to Lysing Matrix E (MP Biomedicals, Santa Ana, USA), snap-frozen in liquid nitrogen, and stored at -80°C for downstream processing. For DNA extractions, samples were homogenized twice by Precellys 24 TissueLyser (6300 rpm/15 sec/10 sec pause; Bertin Technologies, Frankfurt am Main, Germany). Prior to total DNA extraction according to the manufacturer's protocol (FastDNA SPIN kit for Soil; MP Biomedicals, Santa Ana, USA), samples were pretreated with 36 µL Lysozyme (1% w/v in 1% PBS at 37°C for 30 min with intermittent shaking; Merck KGaA, Darmstadt, Germany), 15 µL Proteinase K (>600 mA/mL, Qiagen, Hilden, Germany), and 5 µL Rnase A (100 mg/mL, Qiagen) for 5 min at room temperature. DNA was eluted in nuclease-free water (Merck KGaA, Darmstadt, Germany) and stored at -20°C for MiSeq library preparation.

**16S rRNA gene MiSeq library preparation.** The 16S rRNA gene DNA library for Illumina sequencing was prepared through a 2-step PCR-amplification protocol. In brief, the DNA concentration for each sample was adjusted to 50 ng/µL (NanoDrop 2000c, Thermo Fisher Scientific, Dreieich, Germany). The 16S rDNA regions were PCR-amplified in triplicate (95°C/2 min; 95°C/30 sec, 55°C/30sec, and 72°C/45 sec for 25 cycles; and final extension 72°C/10 min) using the primer set forward 799F (5'-AACMGGATTAGATACCCKG-3') and reverse 1193R (5'-ACGTCATCCCCACCTTCC-3')[12]. To block plant mitochondrial DNA, 10 times the volume 799F or 1193R of the blocking primer (5'-GGCAAGTGTTCTTCGGA/3SpC3/-3') was added to each reaction. Technical replicates were pooled, and leftover primers were enzymatically digested for 30 min at 37°C by adding 3 µL of Exonuclease I (20 U, New England BioLabs, Frankfurt am Main, Germany), 3 µL Antarctic phosphatase (5 U, New England BioLabs, Germany), and 7.32 µL Antarctic phosphatase buffer (New England BioLabs, Germany). The reaction was heat-inactivated (85°C/15 min). Amplicons from the first reaction were PCR-barcoded over 10 cycles (95°C/2 min; 95°C/30 sec, 55°C/30 sec, 72°C/45 sec; final extension 72°C/10 min) using reverse Illumina compatible primers (B1 to B120). PCR reactions were performed using Q5© high-fidelity DNA polymerase (2,000 units/ml, New England Biolabs, Frankfurt am Main, Germany). The PCR replicates from the second reaction were pooled and run through 1.5% (w/v) agarose gel (90 V for ~2.5

h). The amplicons (~0.3 kb) were extracted from gel plugs using QIAquick Gel Extraction Kit (Qiagen, Hilden, Germany) according to the manufacturer's protocol. PCR-products were quantified (NanoDrop 2000c, Thermo Fisher Scientific) and adjusted to 10 ng/µL prior to pooling. The library was cleaned twice with Agencourt AMPure XP Kit (Beckman Coulter, Krefeld, Germany) and submitted for DNA sequencing using the MiSeq Reagent kit v3 with the 2 × 300 bp paired-end sequencing protocol (Illumina Inc., San Diego, USA).

**Sequences processing.** Forward and reverse reads were joined and demultiplexed using Qiime2 pipeline (2018.2.0)[13]. PhiX and chimeric sequences were filtered out using Qiime2-DADA2[14]. All scripts used for read preprocessing are available at https://github.com/hmamine/raw_data_preprocessing. For Shannon and observed OTUs analyses, count reads were rarefied to an even sequencing depth based on the smallest sample size of 601 reads using the R package phyloseq[15]. For community structure analyses, count reads were normalized by cumulative sum scaling normalization factors[16] prior to computing Bray-Curtis distances between samples. To test for significantly enriched OTUs, the data were fitted to a zero-inflated Gaussian mixture model,[16] and OTUs with Benjamini and Hochberg[11] adjusted *P*<.05 were displayed. R scripts used for the described analyses are accessible through https://github.com/hmamine/ZIHJE/community_analysis, and raw data are deposited in http://www.ncbi.nlm.nih.gov/bioproject/549447.

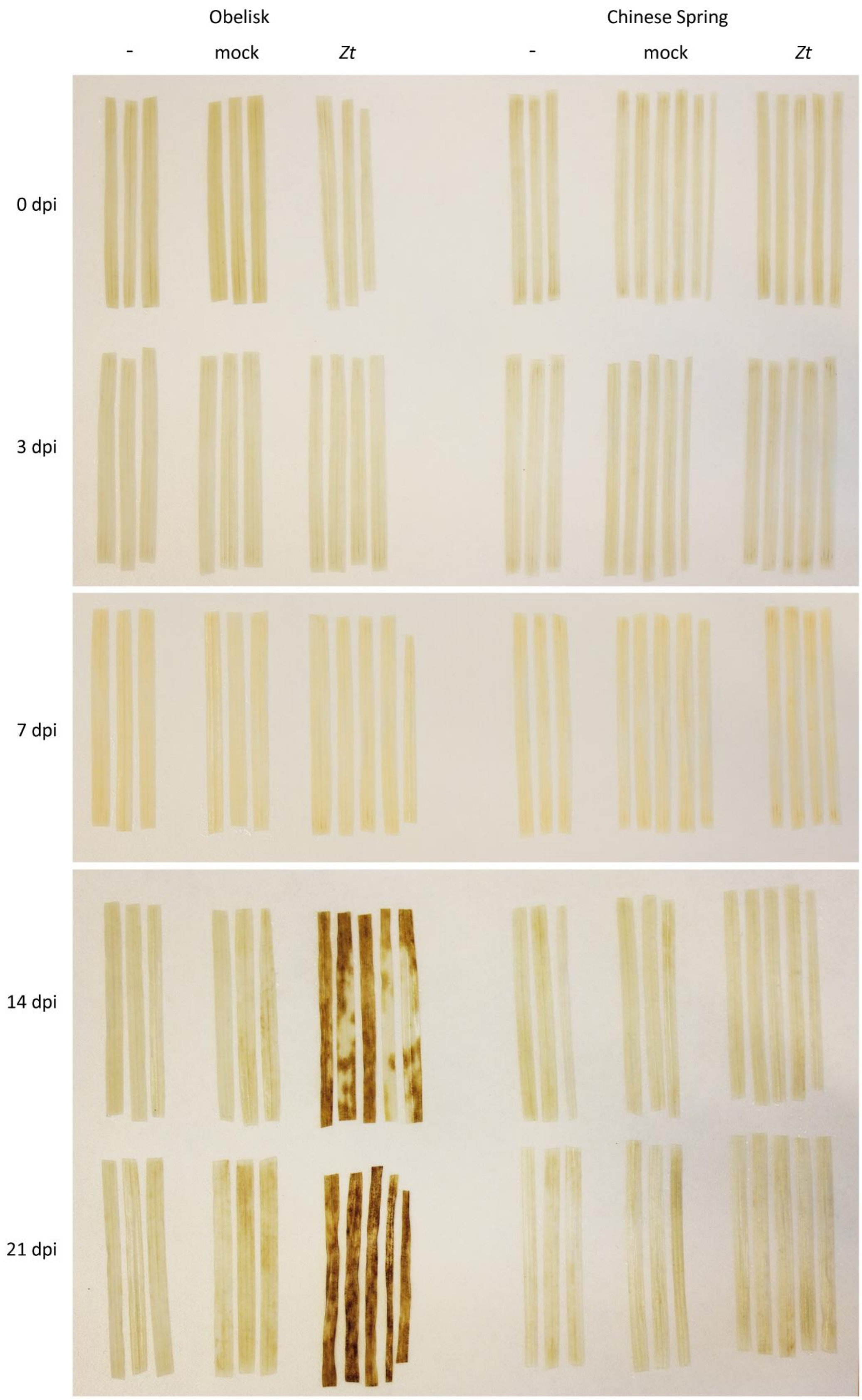


**Fig. S1 | DAB staining after infection with *Z. tritici***

DAB staining of $H_2O_2$ in wheat leaves of cv. Obelisk and cv. Chinese Spring after inoculation with *Z. tritici* (*Zt*) IPO323 at different stages of infection.

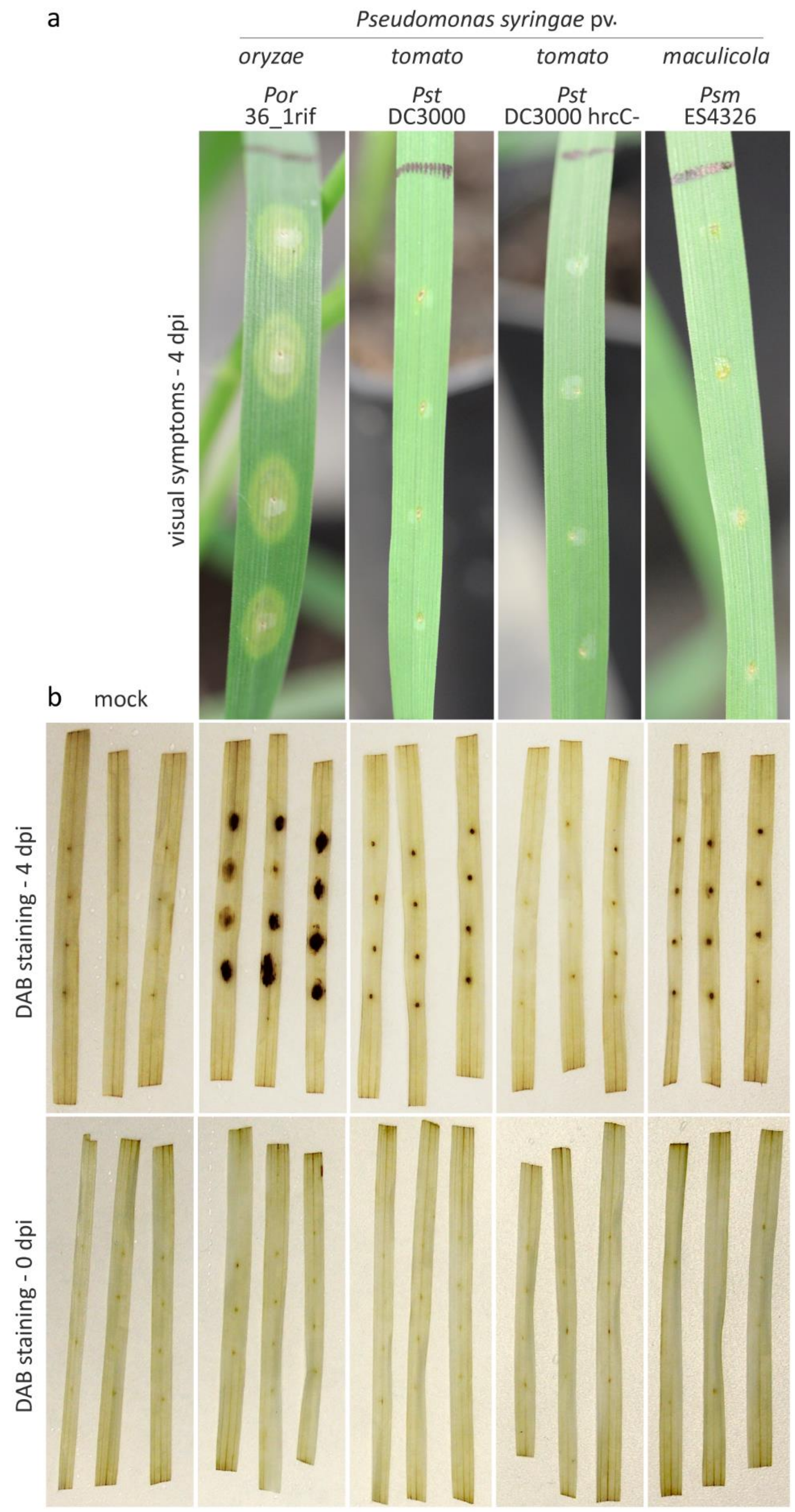


**Fig. S2 | Phenotypes of visible symptoms on wheat infected with *P. syringae* pathovars**

**a**, Visible symptoms of *P. syringae* pv. *oryzae* (*Por*) 36_1rif/ pv. *tomato* (*Pst*) DC3000 and its T3SS mutant hrcC-/ pv. *maculicola* (*Psm*) ES4326 at 4dpi-b on cv. Obelisk, representative symptoms are shown. **b,** DAB staining after infection with *P. syringae* pathovars as in **a** at 0 dpi-b and 4 dpi-b.

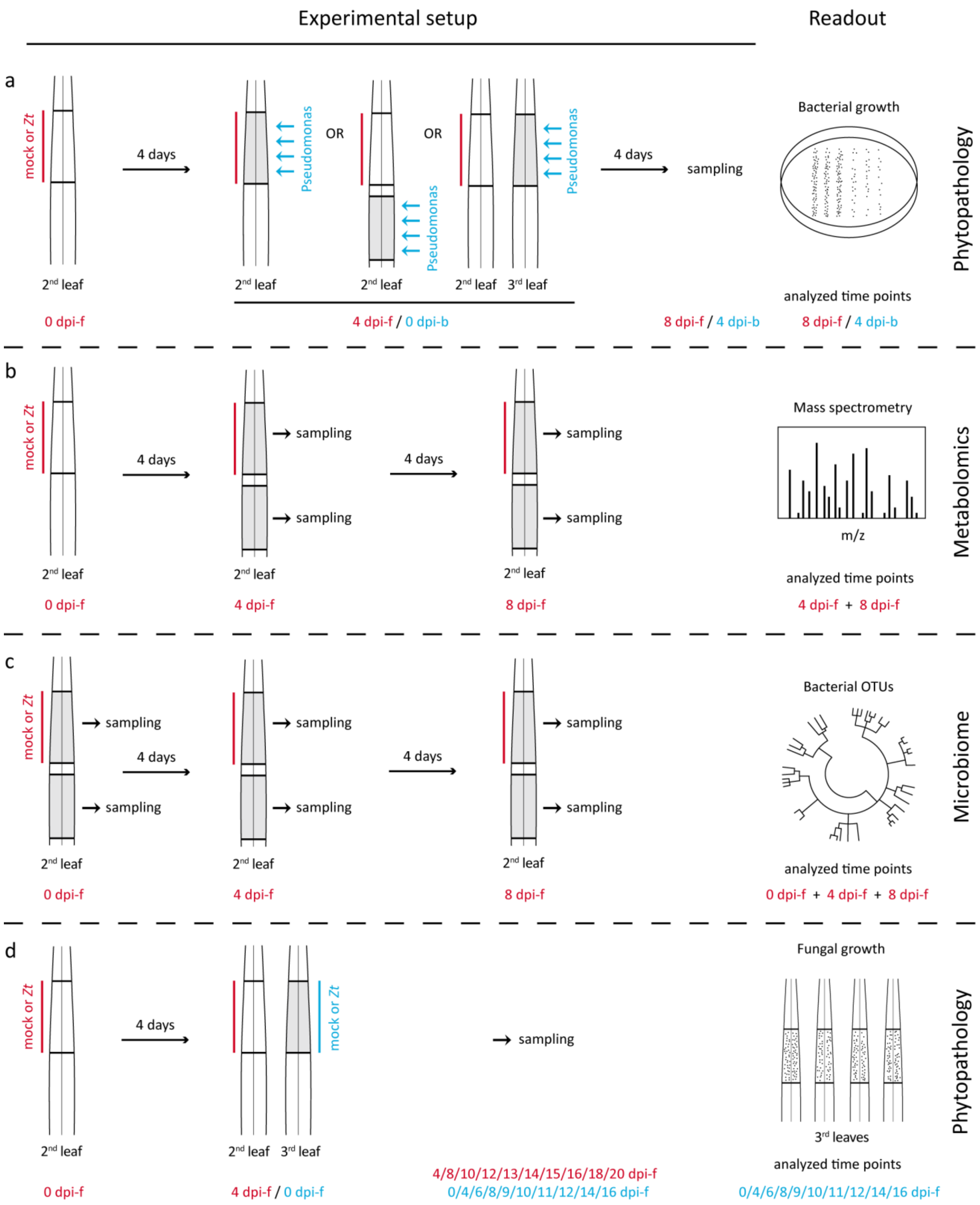


**Fig. S3 | Scheme of the experimental setup and used readouts of the three areas of the study**

**a,** Infection of wheat leaves with *Z. tritici* (*Zt*) IPO323 (red) followed by co-infection with *P. syringae* (blue) at different locations of the leaf/plant at 4 dpi-f. Bacterial performance was monitored at 8 dpi-f/4 dpi-b. **b,** Infection of wheat leaves with *Zt* (red) followed by metabolomics analysis using FT-ICR-MS at 4 and 8 dpi-f in local and adjacent tissue of the leaf. **c,** Infection of wheat leaves with *Zt* (red) followed by analysis of bacterial OTUs at 0, 4 and 8 dpi-f in local and adjacent tissue of the leaf. **d,** Infection of the second leaf with *Zt* (red) followed by systemic co-infection on the third leaf with *Zt* (blue) 4 days after infection of second leaf. Samples for monitoring of fungal growth were taken throughout all infection stages of systemic infection.

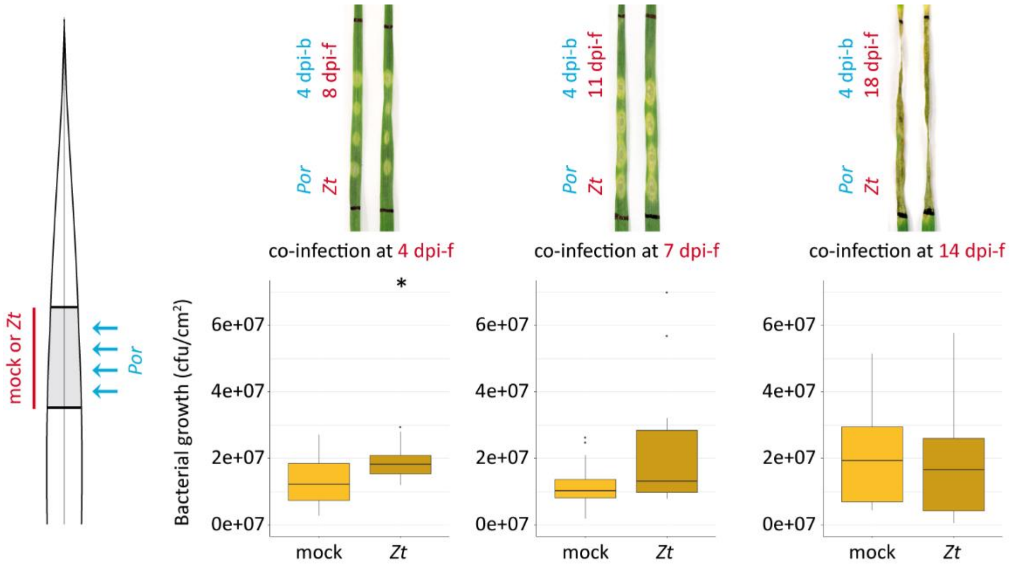


**Fig. S4 | Time course of local co-infection with *Zt* and *Por* covering biotrophic and necrotrophic lifestyles of *Z. tritici***

Local co-infection of cv. Obelisk with *Zt* (red) followed by *Por* (blue) at 4/7/14 dpi-f. Bacterial performance was quantified 4 days after bacterial infection. Statistical analysis was performed using Shapiro-Wilk test of normality followed by Wilcoxon rank-sum test for test of null hypothesis. **P*<.05. Number of biologically independent replicates: 4 dpi-f as in Fig. 2a (n=12), 7 dpi-f mock (n=13), Zt (n=14), 14 dpi-f (n=15).

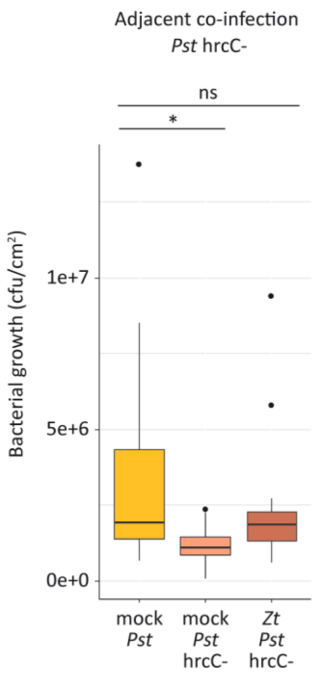


**Fig. S5 | Infection of Obelisk with *Z. tritici* changes replaces T3SS of non-adapted *P. syringae*.**
Adjacent co-infection and quantification of bacterial growth as in Fig. 2c with *Pst* and its T3SS mutant *Pst* hrcC-, respectively. Statistical analysis was performed using Shapiro-Wilk test of normality followed by Wilcoxon rank-sum test for test of null hypothesis. **P*<.05; ns = not significant. Number of biologically independent replicates: *Pst* (n=14), *Pst* hrcC- (n=13).

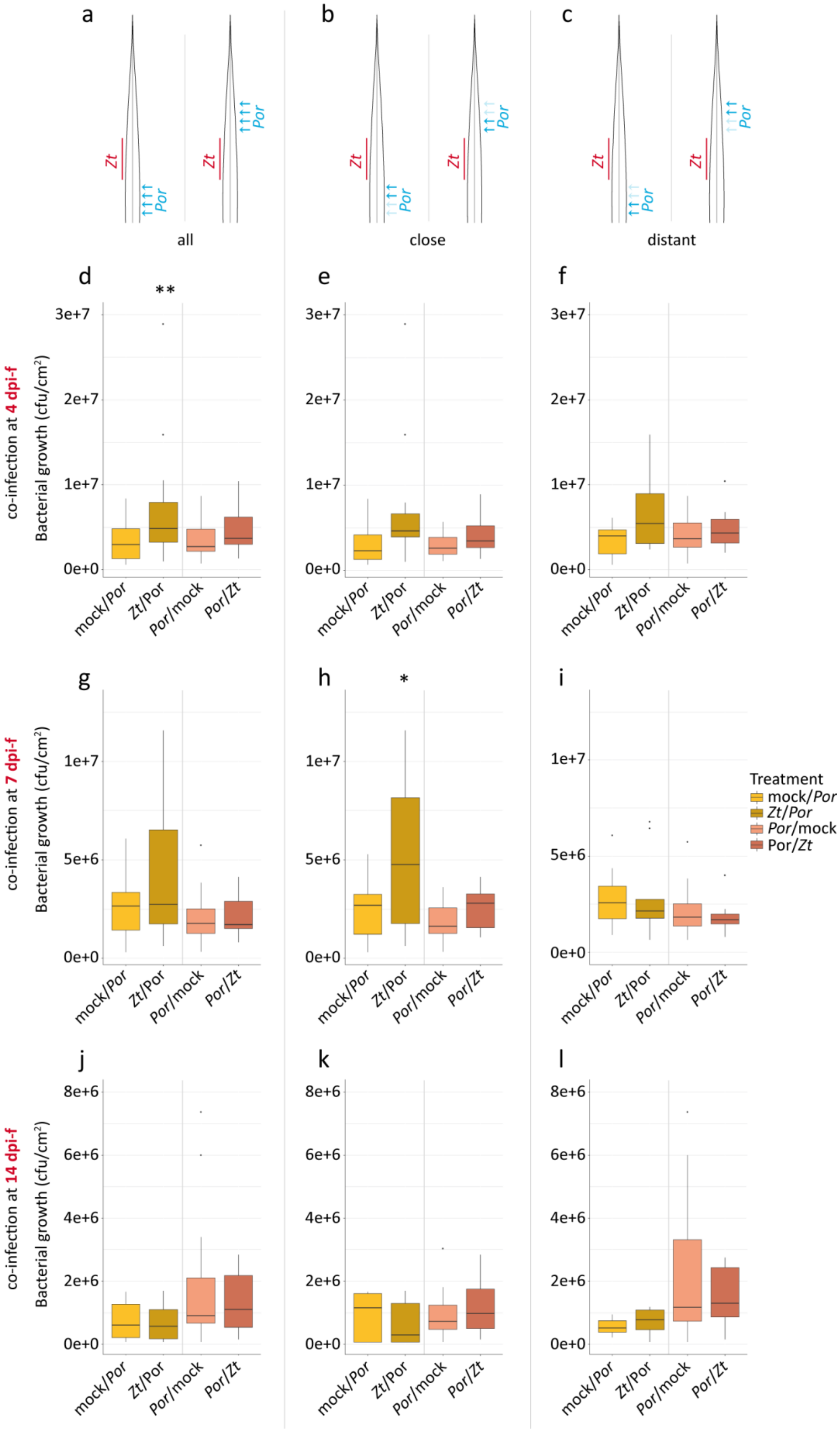
a
b
c
Zt
Por
all
close
distant
d
e
f
g
h
i
j
k
l
**
*
co-infection at 4 dpi-f
co-infection at 7 dpi-f
co-infection at 14 dpi-f
Bacterial growth (cfu/cm²)
3e+7
2e+7
1e+7
0e+0
5e+6
8e+6
6e+6
4e+6
2e+6
mock/Por
Zt/Por
Por/mock
Por/Zt
Treatment
mock/Por
Zt/Por
Por/mock
Por/Zt

275

276

**Fig. S6 | Time course of local infection with *Z. tritici* and adjacent co-infection with Por covering biotrophic and necrotrophic lifestyles of the fungus**

Local infection of cv. Obelisk with *Zt* (red) followed by adjacent co-infection with *Por* (blue) at 4/7/14 dpi-f. Bacterial growth on cv. Obelisk was quantified 4 days after bacterial infection. Bacterial co-infection took place either towards the leaf tip (left panel) or the leaf base (right panel), respectively. **a-c** Schematics of infected leaf area. **d-f** Co-infection with *Por* at 4 dpi-f. **g-i** Co-infection with *Por* at 7 dpi-f. **j-l** Co-infection at 14 dpi-f. **a**,**d**,**g**,**j** Bacterial growth in all samples from adjacent area (combination of close by and distant area). **b**,**e**,**h**,**k** Bacterial growth in samples from area close by to the fungal infection. **c**,**f**,**i**,**l** Bacterial growth in samples from more distant area within the adjacent area. Statistical analysis was performed using Shapiro-Wilk test of normality followed by Wilcoxon rank-sum test for test of null hypothesis. *$P$<.05; **$P$ <.01. Number of biologically independent replicates (n=combined/close by/distant): mock/*Por*: 4 dpi-f 22/11/11; 7 dpi-f 22/11/11; 14 dpi-f 15/9/6. *Zt*/*Por*: 4 dpi-f 21/11/10; 7 dpi-f 21/10/11; 14 dpi-f 16/9/7. *Por*/mock: 4 dpi-f 20/10/10; 7 dpi-f 24/12/12; 14 dpi-f 20/10/10. *Por*/*Zt*: 4 dpi-f 22/11/11; 7 dpi-f 15/8/7; 14 dpi-f 23/14/9.

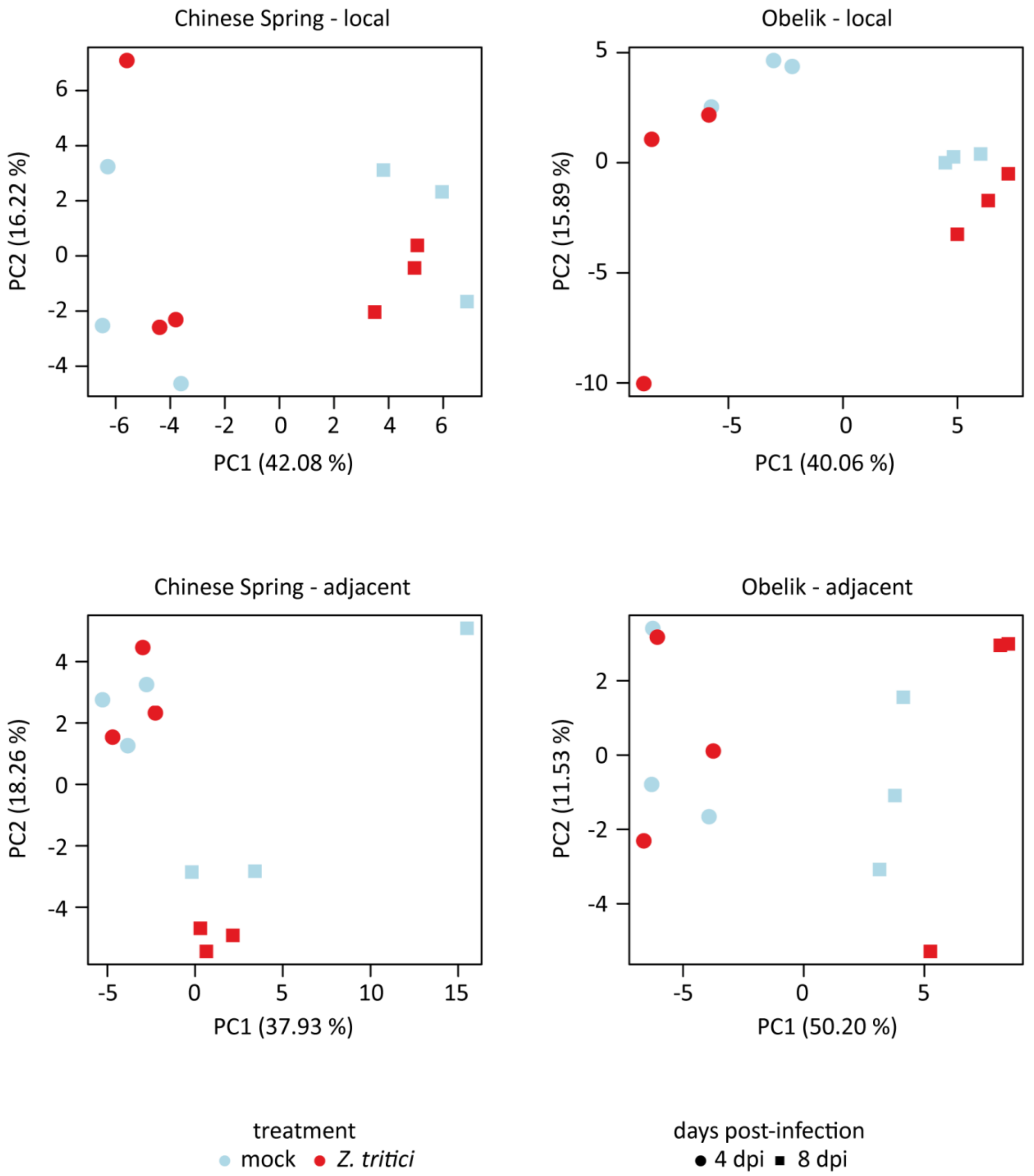


**Fig. S7 | PCA in local and adjacent tissue of *T. aestivum* cultivars Chinese Spring and Obelisk after infection with *Z. tritici***

Principle component analysis (PCA) based on the subset of the metabolomics dataset: **a**, Samples from locally treated Chinese Spring leaf tissue. **b**, Samples from locally treated Obelisk leaf tissue. **c**, Samples from Chinese Spring leaf tissue adjacent to treatment. **d**, Samples from Obelisk leaf tissue adjacent to treatment. Red- and blue-colored shapes designate mock and *Z. tritici*-treated leaf samples. Samples are colored according to treatment; shapes refer to the days after the treatment.

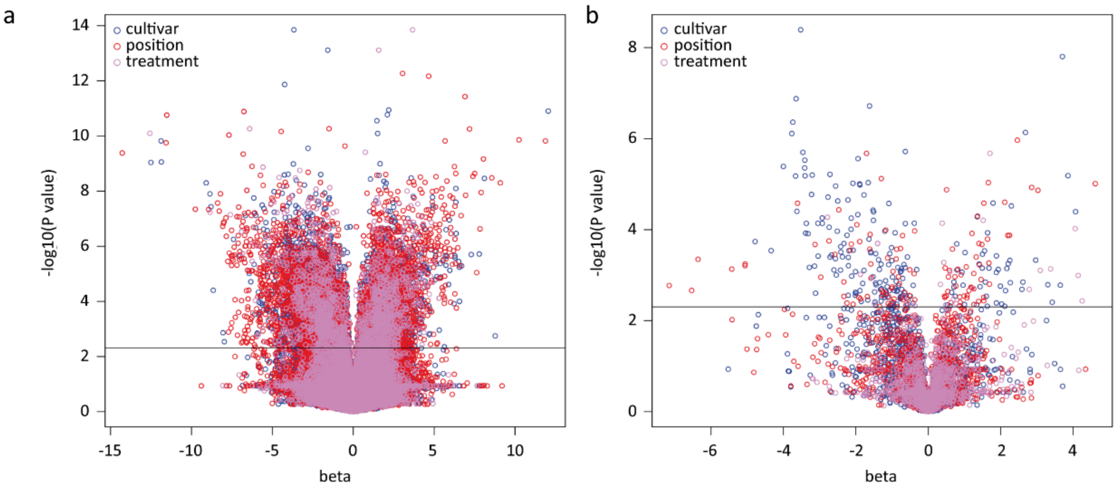


c

| Formula for beta | beta | Set | Conditions | Code |
|---|---|---|---|---|
| log2(**Ob**.mock/**CS**.mock) | negative --> higher in **CS** | cultivar --> CS | mock local 4/8dpi | CS_m-loc_4/8 |
| | | | mock adjacent 4/8dpi | CS_m-adj_4/8 |
| log2(**Ob**.Zt/**CS**.Zt) | | | Zt local 4/8dpi | CS_Zt-loc_4/8 |
| | | | Zt adjacent 4/8dpi | CS_Zt-adj_4/8 |
| log2(**Ob**.mock/**CS**.mock) | positive --> higher in **Ob** | cultivar --> Ob | mock local 4/8dpi | Ob_m-loc_4/8 |
| | | | mock adjacent 4/8dpi | Ob_m-adj_4/8 |
| log2(**Ob**.Zt/**CS**.Zt) | | | Zt local 4/8dpi | Ob_Zt-loc_4/8 |
| | | | Zt adjacent 4/8dpi | Ob_Zt-adj_4/8 |
| log2(CS.**adj**/CS.**loc**) | negative --> higher in **local** | position --> local | CS mock 4/8dpi | loc_CS-m_4/8 |
| | | | CS Zt 4/8dpi | loc_CS-Zt_4/8 |
| log2(Ob.**adj**/Ob.**loc**) | | | Ob mock 4/8dpi | loc_Ob-m_4/8 |
| | | | Ob Zt 4/8dpi | loc_Ob-Zt_4/8 |
| log2(CS.**adj**/CS.**loc**) | positive --> higher in **adjacent** | position --> adjacent | CS mock 4/8dpi | adj_CS-m_4/8 |
| | | | CS Zt 4/8dpi | adj_CS-Zt_4/8 |
| log2(Ob.**adj**/Ob.**loc**) | | | Ob mock 4/8dpi | adj_Ob-m_4/8 |
| | | | Ob Zt 4/8dpi | adj_Ob-Zt_4/8 |
| log2(CS.**Zt**/CS.**mock**) | positive --> higher in **Zt** | treatment --> Zt | CS local 4/8dpi | Zt_CS-loc_4/8 |
| | | | CS adjacent 4/8dpi | Zt_CS-adj_4/8 |
| log2(Ob.**Zt**/Ob.**mock**) | | | Ob local 4/8dpi | Zt_Ob-loc_4/8 |
| | | | Ob adjacent 4/8dpi | Zt_Ob-adj_4/8 |
| log2(CS.**Zt**/CS.**mock**) | negative --> higher in **mock** | treatment --> mock | CS local 4/8dpi | m_CS-loc_4/8 |
| | | | CS adjacent 4/8dpi | m_CS-adj_4/8 |
| log2(Ob.**Zt**/Ob.**mock**) | | | Ob local 4/8dpi | m_Ob-loc_4/8 |
| | | | Ob adjacent 4/8dpi | m_Ob-adj_4/8 |

d

| Compound | Most significant | Additionally significant |
|---|---|---|
| Cinnamyl alcohol | CS_m-loc_4 | CS_m-adj_4, CS_Zt-loc_4, CS_Zt-adj_4, CS_m-loc_8, CS_m-adj_8, CS_Zt-loc_8, CS_Zt-adj_8, adj_CS-m_8 |
| PG(18:3(6Z,9Z,12Z)/0:0) | Ob_Zt-adj_4 | |
| Ethyladipic acid | CS_m-loc_8 | CS_m-loc_4, CS_m-adj_4, CS_Zt-loc_4, CS_Zt-adj_4, CS_m-adj_8, CS_Zt-loc_8, CS_Zt-adj_8 |
| Neocarlinoside | Ob_Zt-loc_4 | Ob_m-loc_4, Ob_m-adj_4, Ob_m-loc_8, Ob_m-adj_8, Ob_Zt-adj_8 |
| Sugeonyl acetate | CS_Zt-adj_8 | CS_m-loc_4, CS_Zt-adj_4, CS_Zt-loc_8, CS_m-adj_8, adj_CS-Zt_8, adj_Ob-m_8 |
| L-Phenylalanine | adj_Ob-m_8 | |
| Diferulic acid | loc_CS-m_4 | CS_m-loc_4 |
| Squalene | adj_CS-Zt_8 | CS_m-loc_4, CS_Zt-adj_4, CS_Zt-adj_8 |
| 4'-O-Methylneobava-iisoflavone 7-O-(2''-p-coumaroylglucoside) | adj_CS-Zt_4 | |
| 3-(2-Carboxyethenyl)-cis-cis-muconate | adj_CS-Zt_4 | Zt_Ob-loc_4 |
| PA(18:3(6Z,9Z,12Z)/0:0) | Zt_CS-loc_4 | CS_Zt-loc_4, loc_CS-Zt_4 |
| 9,10-Dihydroxystearic acid | m_CS-loc_8 | loc_CS-m_4, loc_Ob-m_4, loc_CS-m_8 |
| Phenylpropanolamine | Zt_CS-loc_8 | CS_m-loc_4, CS_Zt-loc_4, CS_m-adj_4, CS_Zt-adj_4, CS_Zt-loc_8, CS_m-adj_8, CS_Zt-adj_8, adj_CS-m_8 |
| Phenyl salicylate | Zt_CS-loc_4 | CS_m-adj_8 |
| Syringic acid | Zt_Ob-adj_8 | loc_CS-Zt_8, CS_m-adj_8 |

**Fig. S8 | Overview metabolomics dataset**

**a,** Vulcano plot of the whole dataset (37664 metabolites), showing significantly accumulating metabolites from the three comparisons cultivar, position and treatment. **b,** Vulcano plot of annotated metabolites showing significant accumulating metabolites from the comparisons as in **a**. **c,** Introduction of the three comparisons and a code for identification of significant conditions. **d,** Metabolites with the lowest *p* value from the three comparisons as in **a**, respectively. Additional significant conditions of the respective metabolites are listed.

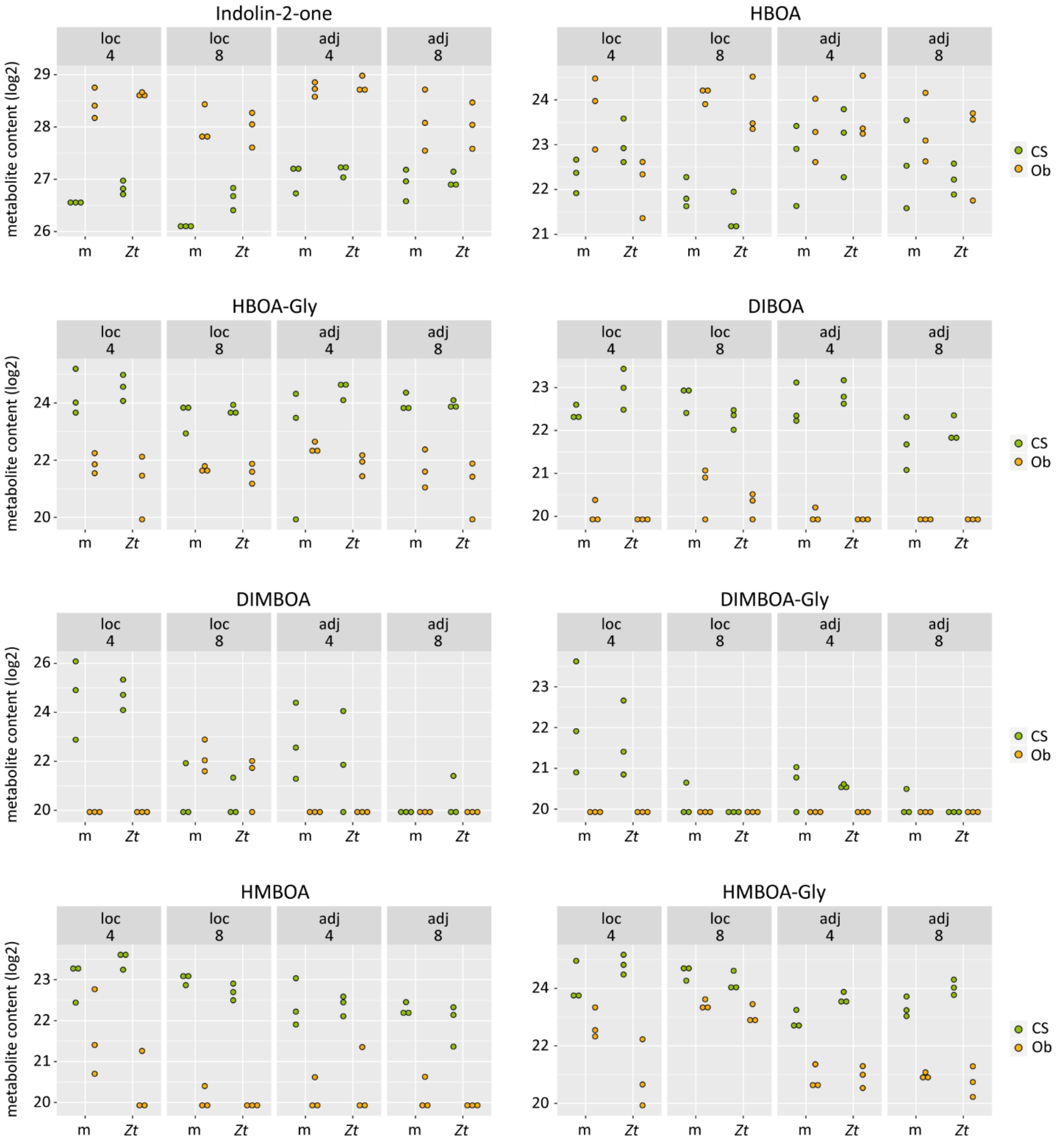


**Fig. S9 | Benzoxazinoids in local and adjacent tissue 4 and 8 days after infection with *Z. tritici***

Detailed view on all compounds from the benzoxazinoid biosynthesis pathway with significant differences as shown in Fig.3. Each dot represents one independent biological sample.

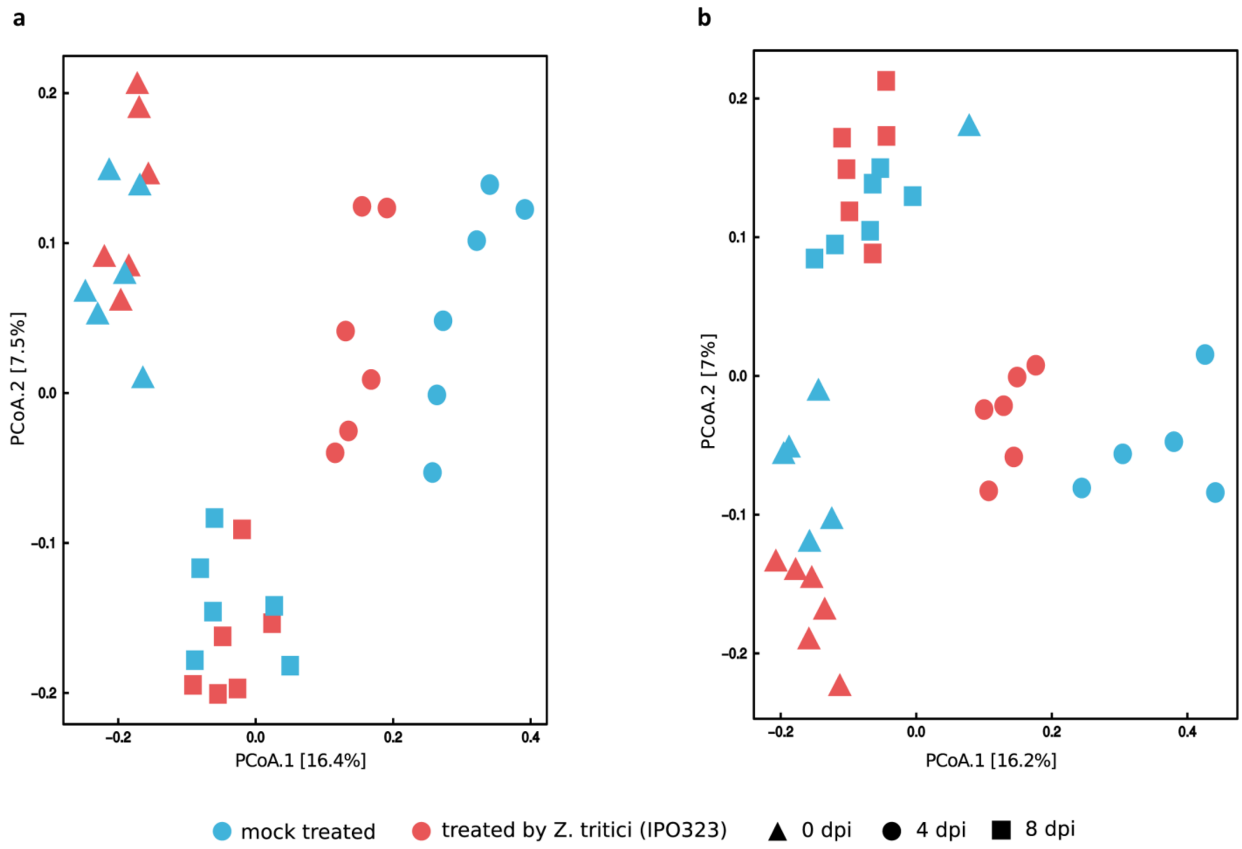


**Fig. S10 | Community structure of the leaf microbiota upon *Z. tritici* treatment in *T. aestivum* cultivar Obelisk.**

**a** and **b** indicate principal coordinates analysis computed on Bray-Curtis (BC) distances of local (**a**) and adjacent (**b**) leaf tissues, respectively. Each shape in the plot represents the community structure of one sample. Red- and blue-colored shapes designate mock and *Z. tritici*-treated leaf samples. Prior to compute the BC distances between samples, reads were normalized by cumulative sum scaling normalization factors.

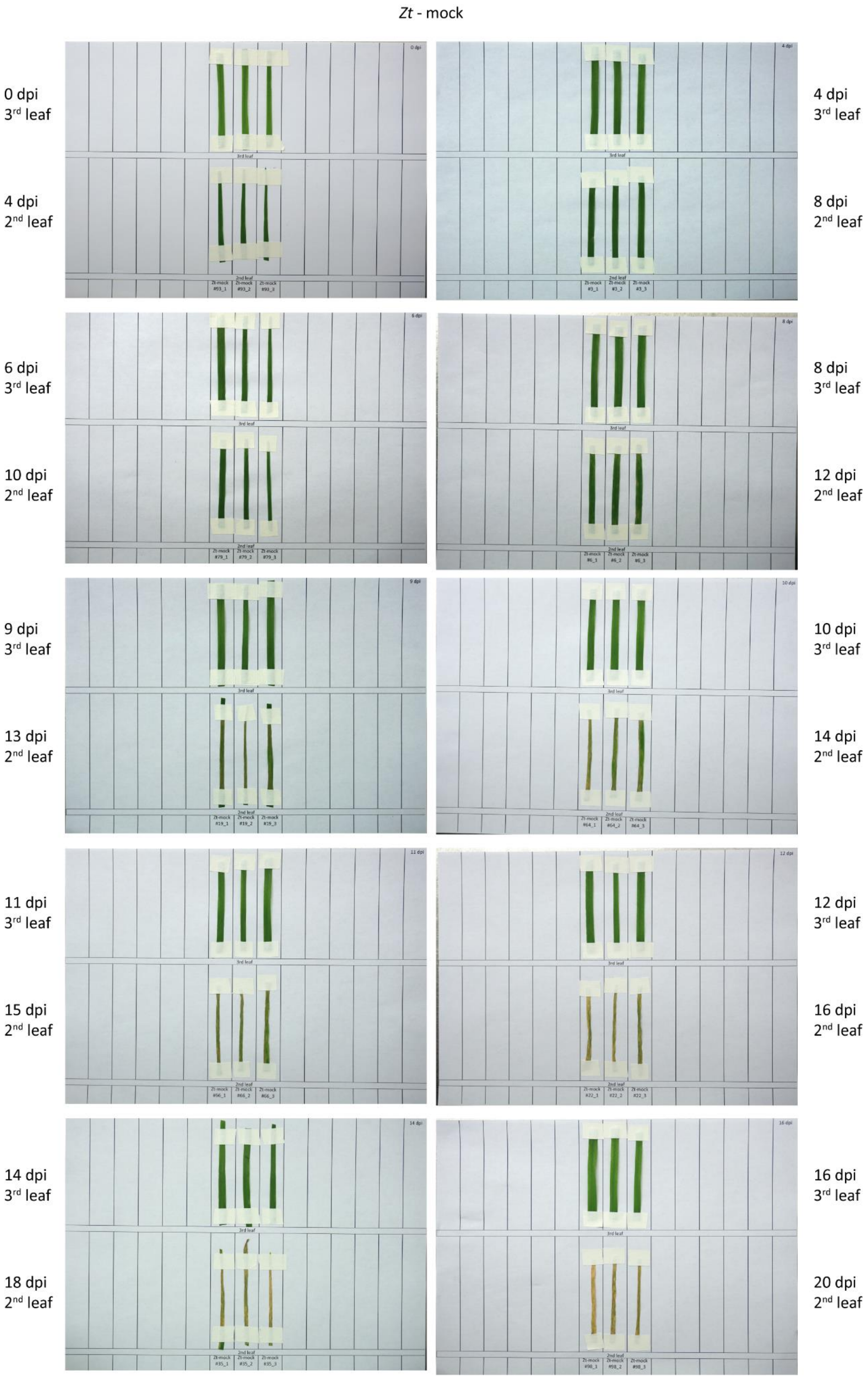


**Fig. S11 | Documentation of symptom development during fungal co-infection**

Visual phenotypes of leaves treated with *Zt* (second leaf)-mock (third leaf) used in Fig.6.

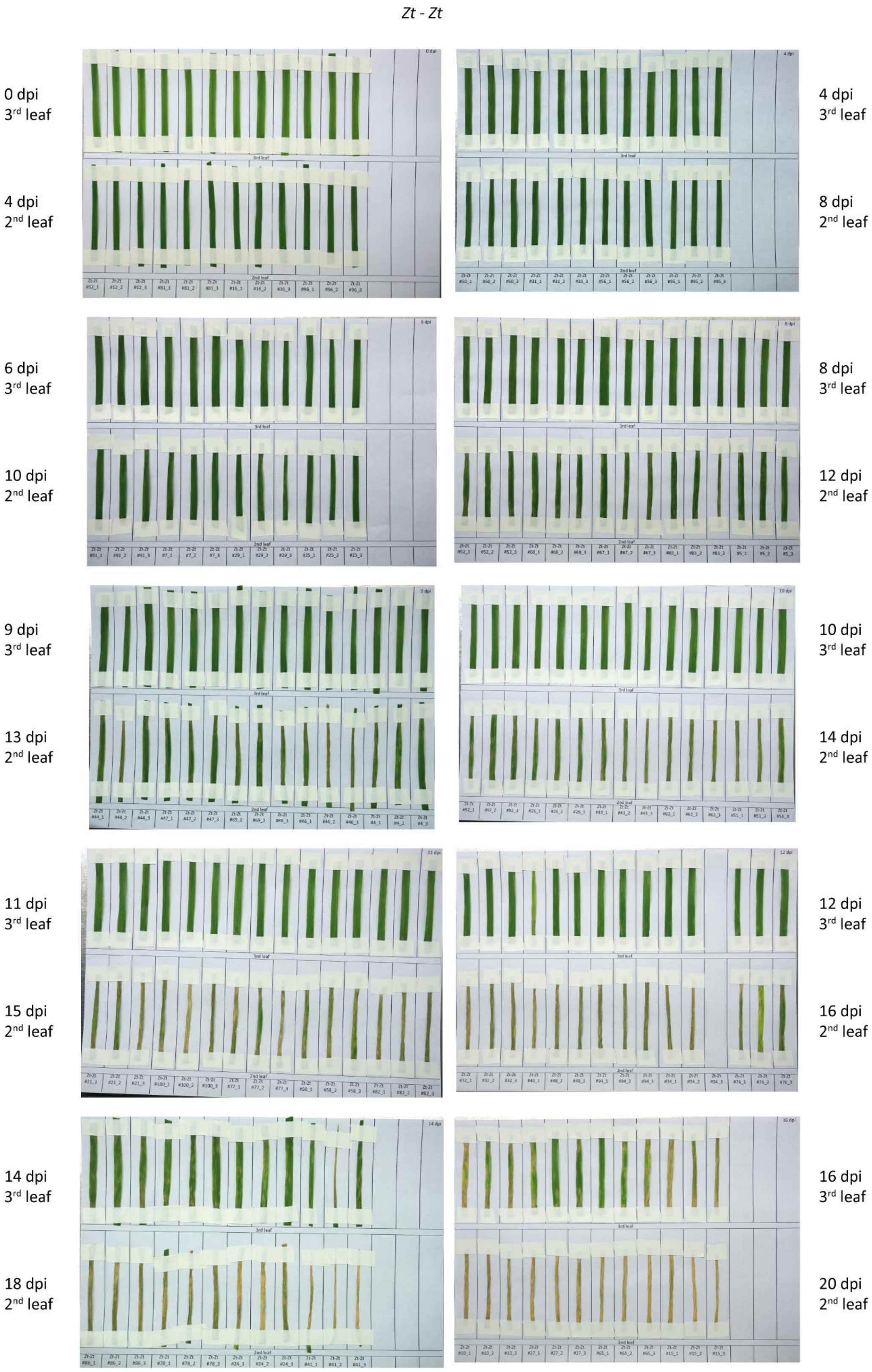


**Fig. S12 | Documentation of symptom development during fungal co-infection**

Visual phenotypes of leaves treated with *Zt* (second leaf)-*Zt* (third leaf) used in Fig.6.

mock - *Zt*

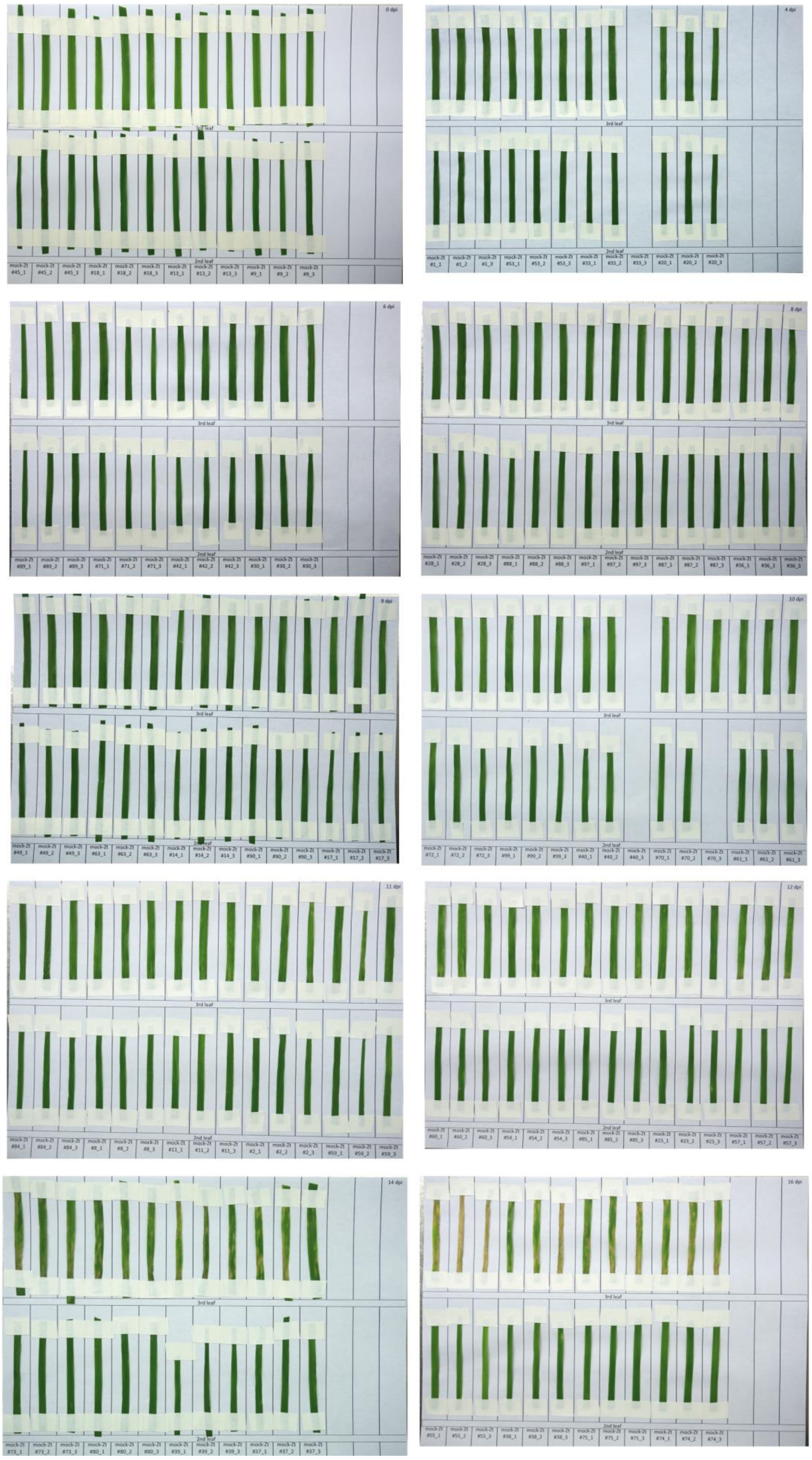


**Fig. S13 | Documentation of symptom development during fungal co-infection**

Visual phenotypes of leaves treated with mock (second leaf)-*Zt* (third leaf) used in Fig.6.

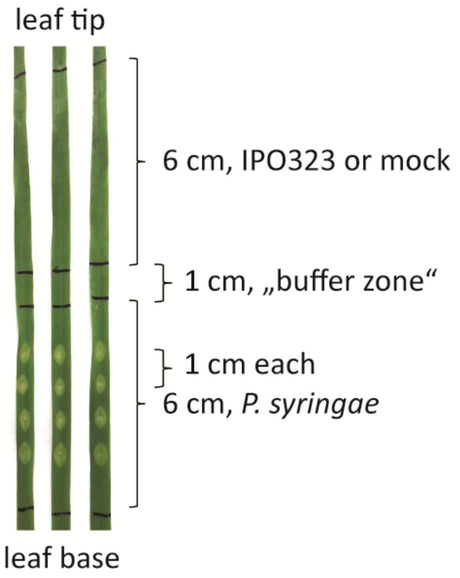


**Fig. S14 | Schematic of adjacent bacterial co-infection**

**Table S3 | Fisher’s exact test of differentially accumulating metabolites**

### Results Fisher’s exact test

| | cultivar | position | treatment |
|---|---|---|---|
| Ob vs CS | p-value < 2.2e-16 | | |
| local (Ob vs CS) | | p-value < 2.2e-16 | |
| adj (Ob vs CS) | | p-value < 2.2e-16 | |
| Zt up (Ob vs CS) | | | p-value < 2.2e-16 |
| Zt down (Ob vs CS) | | | p-value = 0.00578 |

### Is there a significant difference in the number of significantly differential accumulating metabolites in Ob and CS? (set: cultivar)

```
       sig     ns
Ob_up 3820 297482
CS_up 5191 296031

Fisher's Exact Test for Count Data:
data:  x
p-value < 2.2e-16
alternative hypothesis: true odds ratio is not equal to 1
95 percent confidence interval:
 0.7019548 0.7639351
sample estimates:
odds ratio
  0.732303
```

### Is there a significant difference between Ob and CS in local tissue? (set: position)

```
           sig    ns
Ob_up loc 2466 27914
CS_up loc 4465 29906

Fisher's Exact Test for Count Data:
data:  x
p-value < 2.2e-16
alternative hypothesis: true odds ratio is not equal to 1
95 percent confidence interval:
 0.5615801 0.6233487
sample estimates:
odds ratio
 0.5917177
```

```
# Is there also a significant difference between Ob and CS in adjacent tissue? (set: position)

            sig    ns
Ob_up adj 2474 30945
CS_up adj 3390 27374

Fisher's Exact Test for Count Data:
data:  x
p-value < 2.2e-16
alternative hypothesis: true odds ratio is not equal to 1
95 percent confidence interval:
 0.6111900 0.6818176
sample estimates:
odds ratio
 0.6455889


# Is there a significant difference between Ob and CS in the number of metabolites upregulated by Zt treatment? (set: treatment)

          sig    ns
Ob_Zt up 397 28248
CS_Zt up 839 30063

Fisher's Exact Test for Count Data:
data:  x
p-value < 2.2e-16
alternative hypothesis: true odds ratio is not equal to 1
95 percent confidence interval:
 0.4452784 0.5687844
sample estimates:
odds ratio
 0.5035934


# Is there a significant difference between Ob and CS in the number of metabolites downregulated by Zt treatment? (set: treatment)

            sig    ns
Ob_Zt down 584 29060
CS_Zt down 584 24672

Fisher's Exact Test for Count Data:
data:  x
p-value = 0.00578
alternative hypothesis: true odds ratio is not equal to 1
95 percent confidence interval:
 0.7547104 0.9550504
sample estimates:
odds ratio
 0.8490065
```

**Table S4 | perMANOVA test**

| | *T. aestivum* cultivar Chinese Spring | | | | | |
|---|---|---|---|---|---|---|
| | 0 dpi | | 4 dpi | | 8 dpi | |
| | local | adjacent | local | adjacent | local | adjacent |
| explain. var. | 0.100 | 0.185 | 0.264 | 0.141 | 0.177 | 0.135 |
| residu. var. | 0.899 | 0.8141 | 0.735 | 0.858 | 0.822 | 0864 |
| *p-value* | 0.166 | 0.003 | 0.003 | 0.017 | 0.003 | 0.032 |

| | *T. aestivum* cultivar Obelisk | | | | | |
|---|---|---|---|---|---|---|
| | 0 dpi | | 4 dpi | | 8 dpi | |
| | local | adjacent | local | adjacent | local | adjacent |
| explain. var. | 0.118 | 0.135 | 0.178 | 0.193 | 0.096 | 0.100 |
| residu. var. | 0.881 | 0.864 | 0.821 | 0.806 | 0.903 | 0.899 |
| *p-value* | 0.003 | 0.002 | 0.002 | 0.004 | 0.257 | 0.168 |

explain. var. : explained variance      residu. var. : residuals variance      dpi : days post-infection

**Table S5 | KEGG pathways**

| | KEGG name (Reference pathway) | KEGG entry |
|---|---|---|
| I | Biosynthesis of plant secondary metabolites | map01060 |
| II | Biosynthesis of plant hormones | map01070 |
| III | Jasmonic acid biosynthesis | M00113 |
| IV | Abscisic acid biosynthesis | M00372 |
| V | Betaine biosynthesis | M00555 |
| VI | Monolignol biosynthesis | M00039 |
| VII | Flavonoid biosynthesis | map00941 |
| VIII | Flavanone biosynthesis | M00137 |
| IX | Phenylpropanoid biosynthesis | map00940 |
| X | Polyamine biosynthesis | M00133 and M00134 |
| XI | Ascorbate biosynthesis, plants | M00114 |
| XII | Shikimate pathway | M00022 |
| XIII | alpha-Linolenic acid metabolism | map00592 |
| XIV | Phenylalanine metabolism | map00360 |
| XV | Benzoxazinoid biosynthesis | map00402 |
