## Supplemental Table 1 for "Hemibiotrophic fungal pathogen induces systemic susceptibility and systemic shifts in wheat metabolome and microbiome composition"

| Compound name | Molecular formula | KEGG ID |
|---|---|---|
| (-)-Jolkinol A | C29H36O6 | |
| (-)-Jolkinol B | C29H36O5 | |
| (+)-Gallocatechin | C15H14O7 | C12127 |
| (9R,13R)-1a,1b-dihomo-jasmonic acid | C14H22O3 | |
| (9Z)-Octadecenoic acid | C18H34O2 | |
| 11R-octadecanoyloxyoctadeca-9Z,12Z,15Z- trienoic acid | C36H64O4 | |
| 12-hydroxy-Jasmonic acid | C12H18O4 | |
| 12-Hydroxy-jasmonic acid-Ile | C18H29NO5 | |
| 12-OPDA | C18H28O3 | C01226 |
| 1-Methoxyficifolinol | C26H30O5 | |
| 1-O-Galloyl-beta-D-glucose | C13H16O10 | C01158 |
| 1-O-Sinapoyl-beta-D-glucose | C17H22O10 | C01175 |
| 2,5-Dimethoxystilbene | C16H16O2 | C15131 |
| 2C-Methyl-D-erythritol-4-phosphate | C5H13O7P | C11434 |
| 2-Dehydro-3-deoxy-D-arabino-heptonate | C7H13O10P | C04691 |
| 2-Hydroxy-4-methoxybenzophenone | C14H12O3 | C14285 |
| 2-Oxoglutarate | C5H6O5 | C00026 |
| 2-Phenylacetamide | C8H9NO | |
| 3-(2-Carboxyethenyl)-cis,cis-muconate | C9H8O6 | |
| 3, 5-Dinitrosalicylic acid | C7H4N2O7 | |
| 3-Amino-4, 7-dihydroxycoumarin | C9H7NO4 | C12468 |
| 3-Hydroxyadipic acid | C6H10O5 | C02360 |
| 3-Hydroxyindolin-2-one | C8H7NO2 | C11130 |
| 3-Phospho-D-glycerate | C3H7O7P | C00197 |
| 4, 4'-dihydroxy-3, 5-dimethoxydihydrostilbene | C16H18O4 | C10256 |
| 4, 4'-Dihydroxystilbene | C14H12O2 | C14233 |
| 4,2,3',4'-Tetrahydroxychalcone 4'-O-(2”-O-p-coumaroyl) glucoside | C30H28O12 | |
| 4-Hydroxycinnamyl alcohol 4-D-glucoside | C15H20O7 | C05855 |
| 4-Hydroxynonenal | C9H16O2 | C21642 |
| 4-Methoxycinnamic acid | C10H10O3 | |
| 4-Methylumbelliferyl acetate | C12H10O4 | C03837 |
| 4'-O-Methylneobavaisoflavone 7-O-(2''-p-coumaroylglucoside) | C36H36O11 | |
| 5'-Butyrylphosphoinosine | C14H19N4O9P | C06435 |
| 5-Hydroxy-6,7,3',4',5'-pentamethoxyflavanone 5-O-rhamnoside | C26H32O12 | |
| 5-Hydroxyconiferyl alcohol | C10H12O4 | C12205 |
| 5-Hydroxyferulate | C10H10O5 | C05619 |
| 5-Methoxypodophyllotoxin | C23H24O9 | |
| 5-O-Methyleriodictyol 7-glucosyl-(1, 4)-galactoside | C28H34O16 | |
| 6,9,12,15-octadecatetraenoic acid | C18H28O2 | C16300 |
| 6-Hydroxykaempferol 3,5,7-trimethyl ether | C18H16O7 | |
| 7-Heptadecynoic acid | C17H30O2 | |
| 8, 1R, 2R, 3, Oxo, 2, Z, pent, 2, enyl}cyclopentyloctanoate | C18H30O3 | C04780 |
| 8,3',4'-Trihydroxy-5,7-dimethoxy-4-phenylcoumarin | C17H14O7 | |
| 8-Methoxy-13-hydroxy-9,11-octadecadienoic acid | C19H34O4 | |
| 9,10-Dihydroxystearic acid | C18H34O6 | |
| 9-cis-Retinol | C20H30O | C16682 |
| 9-Hydroperoxy-10E,12,15Z-octadecatrienoic acid | C18H30O4 | |
| 9-Hydroxy-5Z-nonenoic acid | C9H16O3 | |
| 9S,10S,11R-trihydroxy-12Z,15Z-octadecadienoic acid | C18H32O5 | |
| 9S,10S,11R-trihydroxy-12Z-octadecenoic acid | C18H34O5 | |
| 9Z-Octadecenedioic acid | C18H32O4 | |
| Abietic acid | C20H30O2 | C06087 |
| Abscisic acid | C15H20O4 | C06082 |
| Abscisic alcohol | C15H22O3 | C13456 |
| Abscisic aldehyde | C15H20O3 | C13455 |
| Acetyl-CoA | C23H38N7O17P3S | C00024 |
| all trans Phytoene | C40H64 | C05413 |
| alpha,alpha'-diethyl-3, 4, 4'-stilbenetriol | C18H20O3 | C14740 |
| alpha-D-Glucose | C6H12O6 | C00267 |
| alpha-D-Glucose-6-phosphate | C6H13O9P | C00668 |
| Alpha-Linolenic acid | C18H30O2 | C06427 |
| alpha-Tocopherol | C29H50O2 | C02477 |

| | | |
|---|---|---|
| alpha-Zearalenol | C18H24O5 | C14750 |
| Aminoadipic acid | C6H11NO4 | C00956 |
| Aminocyclopropanecarboxylate | C4H7NO2 | C01234 |
| AMP | C10H14N5O7P | C00020 |
| Antheraxanthin | C40H56O3 | C08579 |
| Ascorbate | C6H8O6 | C00072 |
| Aureusidin 6-O-glucoside | C21H20O11 | |
| Avenoleic acid | C18H32O3 | |
| Azaleatin | C16H12O7 | C10022 |
| Azelaic acid | C9H16O4 | C08261 |
| Benzimidazole | C7H6N2 | |
| beta Carotene | C40H56 | C02094 |
| Betaine | C5H11NO2 | C00719 |
| Brassinolide | C28H48O6 | C08814 |
| Caffeate | C9H8O4 | C01197 |
| Caffeyl alcohol | C9H10O3 | C12206 |
| Campesterol | C28H48O | C01789 |
| Capsidiol | C15H24O2 | C09627 |
| Chlorogenic acid | C16H18O9 | C00852 |
| Chorismate | C10H10O6 | C00251 |
| Chrysanthemic acid | C10H16O2 | C09842 |
| Cinnamoyltyramine | C17H17N1O2 | |
| Cinnamyl alcohol | C9H10O | C02394 |
| cis-Aconitate | C6H6O6 | C00417 |
| cis-Stilbene oxide | C14H12O | C16014 |
| Citrate | C6H8O7 | C00158 |
| Coniferyl acetate | C12H14O4 | |
| Coumaroylhydroxyagmatine | C14H20N4O3 | |
| D-Alpha-aminobutyric acid | C4H9NO2 | C02261 |
| Dehydroabietic acid | C20H28O2 | C12078 |
| Dehydroabietinal | C20H28O | |
| Dehydrospermidine | C7H17N3 | C15853 |
| de-Hypoxanthine futalosine | C14H16O7 | C17010 |
| Deoxy-D-xylulose-5-phosphate | C5H11O7P | C11437 |
| D-Erythrose | C4H9O7P | C00279 |
| D-Glyceraldehyde phosphate | C3H7O6P | C00118 |
| DIBOA | C8H7NO4 | C15770 |
| DIBOA-Gly | C14H17NO9 | C15772 |
| Diferulic acid | C20H18O8 | C10446 |
| Dihydroconiferyl alcohol | C10H14O3 | C10448 |
| Dihydrokaempferol | C15H12O6 | C00974 |
| Dihydropinosylvin | C14H14O2 | C10254 |
| Dihydroresveratrol | C14H14O3 | C10255 |
| DIM2BOA | C10H11NO6 | |
| DIM2BOA-Gly | C16H21NO11 | |
| DIMBOA | C9H9NO5 | C04720 |
| DIMBOA-Gly | C15H19NO10 | C04831 |
| D-Ribitol 5-phosphate | C5H13O8P | C01068 |
| Ethyladipic acid | C8H14O4 | |
| Ethylene | C2H4 | C06547 |
| Fenpicoxamid | C31H38N2O11 | |
| Ferulasäure | C10H10O4 | C01494 |
| Feruloylagmatine | C15H22N4O3 | C18325 |
| Feruloylhydroxyagmatine | C15H22N4O4 | |
| Feruloylputrescine | C14H20N2O3 | C10497 |
| Ficifolinol | C25H28O4 | |
| Fumarate | C4H4O4 | C00122 |
| Geranyl diphosphate | C10H20O7P2 | C00341 |
| Geranyl formate | C11H18O2 | C12294 |
| Geranylgeranyl diphosphate | C20H36O7P2 | C00353 |
| Geranylhydroquinone | C16H22O2 | C10793 |
| Gibberellin | C19H24O6 | C00859 |
| Gibberellin A12 aldehyde | C20H28O3 | C06093 |

| | | |
|---|---|---|
| Gingerglycolipid A | C33H56O14 | |
| Glutathione | C10H17N3O6S | C00051 |
| HBOA | C8H7NO3 | C15769 |
| HBOA-Gly | C14H17NO8 | |
| HDM2BOA | C11H13NO6 | |
| HDM2BOA-Gly | C17H23NO11 | |
| HDMBOA | C10H11NO5 | |
| HDMBOA-Gly | C16H21NO8 | |
| HMBOA | C9H9NO4 | |
| HMBOA-Gly | C15H19NO9 | |
| Hordatine A | C28H38N8O4 | C08307 |
| Hordatine A + 2 hex | C40H58N8O14 | |
| hordatine A + hex | C34H48N8O9 | |
| Hordatine B | C29H40N8O5 | C08308 |
| Hordatine B + 2 hex | C41H60N8O15 | |
| Hordatine B + hex | C35H50N8O10 | |
| Hordatine C | C30H42N8O6 | |
| Hordatine C + 2 hex | C42H62N8O16 | |
| Hordatine C + hex | C36H52N8O11 | |
| Hordatine D | C31H44N8O7 | |
| Hordatine D + 2 hex | C43H64N8O17 | |
| Hordatine D + hex | C37H54N8O12 | |
| Hydroxy-hordatine A | C28H38N8O5 | |
| Hydroxy-hordatine A + 2 hex | C40H50N8O16 | |
| Hydroxy-hordatine A + hex | C34H48N8O10 | |
| Hydroxy-hordatine B | C29H40N8O6 | |
| Hydroxy-hordatine B + hex | C35H50N8O11 | |
| Hydroxy-hordatine C | C30H42N8O7 | |
| Hydroxy-hordatine C + hex | C36H52N8O12 | |
| Hydroxy-hordatine D | C31H44N8O8 | |
| Hydroxy-hordatine D + 2 hex | C43H64N8O18 | |
| Hydroxy-hordatine D + hex | C37H54N8O13 | |
| IMP | C10H13N4O8P | C00130 |
| Indole | C8H7N | C00463 |
| Indoleacetic acid | C10H9NO2 | C00954 |
| Indoleglycerol phosphate | C11H14NO6P | C03506 |
| Indolin-2-one | C8H7NO | C12312 |
| Isopentenyl | C5H12O7P2 | C00129 |
| Jasmonic acid | C12H18O3 | C08491 |
| Jasmonic acid isloeucin | C18H29NO4 | |
| Kaempferol 3-(2”-p-coumaryl-alpha-L-arabinopyranoside) | C29H24O12 | |
| Kaempferol 3-(4''-p-coumaroylglucoside) | C30H26O13 | |
| Kaempferol 3,5-dimethyl ether | C17H14O6 | |
| Kaempferol 3-rhamnoside-(1->2)-rhamnoside | C27H30O14 | |
| Kaempferol 7-rhamnoside | C21H20O10 | |
| Kaempferol-3-glucoside-3-rhamnoside | C27H30O15 | |
| Kaurene | C20H32 | C06090 |
| L-Alanine | C3H7NO2 | C00041 |
| L-Arginine | C6H14N4O2 | C00062 |
| L-Asparagine | C4H8N2O3 | |
| L-Aspartate | C4H7NO4 | C00049 |
| L-Galactono-1,4-lactone | C6H10O6 | |
| L-Glutamine | C5H10N2O3 | C00064 |
| L-Histidine | C6H9N3O2 | C00135 |
| Lignoceric acid | C24H48O2 | C08320 |
| Linoelaidic acid | C18H32O2 | |
| L-Leucine | C6H13NO2 | C00123 |
| L-Lysine | C6H14N2O2 | C00047 |
| L-Malate | C4H6O5 | C00149 |
| L-Methionine | C5H11NO2S | C00073 |
| L-Phenylalanine | C9H11NO2 | C00079 |
| L-Proline | C5H9NO2 | C00148 |
| L-Selenocysteine | C3H7NO2Se | C05688 |

| | | |
|---|---|---|
| L-Threonine | C4H9NO3 | C00188 |
| L-Tryptophan | C11H12N2O2 | C00078 |
| L-Tyrosine | C9H11NO3 | C00082 |
| Malonyl-CoA | C24H38N7O19P3S | C00083 |
| Malvidin | C17H15ClO7 | C08716 |
| MBOA | C8H7O3N | |
| Methyl 9,12-dihydroxy-13-oxo-10-octadecenoate | C19H34O5 | |
| Methyljasmonate | C13H20O3 | C11512 |
| Methylsalicylate | C8H8O3 | C12305 |
| Mevalonate | C6H12O4 | C00418 |
| Myristic acid | C14H28O2 | C06424 |
| N-(4-Guanidinobutyl)-4-hydroxycinnamide | C14H20N4O2 | C04498 |
| Naringenin | C15H12O5 | C00509 |
| Naringenin 7-O-beta-D-glucoside | C21H22O10 | C09099 |
| N-Caffeoyl serotonin | C19H18N2O4 | |
| N-Caffeoylputrescine | C13H18N2O3 | C03002 |
| N-Coumaroyl serotonin | C19H18N2O3 | |
| Neocarlinoside | C26H28O15 | C10109 |
| N-Feruloyl serotonin | C20H20N2O4 | |
| Niacinamide | C6H6N2O | C00153 |
| O-Feruloylquinate | C17H20O9 | C02572 |
| Oxaloacetate | C4H4O5 | C00036 |
| PA(18:3(6Z,9Z,12Z)/0:0) | C21H37O7P | |
| Palmitoyl-CoA | C37H66N7O17P3S | C00154 |
| p-Coumaroyl-D-glucose | C15H18O8 | C16827 |
| PE(18:3(6Z,9Z,12Z)/0:0) | C23H42NO7P | |
| Pedilstatin | C30H40O7 | C09143 |
| PG(16:1(9Z)/0:0) | C22H43O9P | |
| PG(18:3(6Z,9Z,12Z)/0:0) | C24H43O9P | |
| PG(8,0,8,0) | C22H43O10P | |
| Phenyl salicylate | C13H10O3 | C14163 |
| Phenylpropanolamine | C9H13NO | C16719 |
| Phenylpropiolic acid | C9H6O2 | |
| Phorbol | C20H28O6 | C09155 |
| Phospho alpha D-ribose | C5H13O14P3 | C00119 |
| Phosphoenolpyruvate | C3H5O6P | C00074 |
| Phthalylamide | C8H6NO3 | C06374 |
| Phyllocoumarin | C18H14O7 | |
| PI(0:0/10:5) | C9H17O11PHC10H9O | |
| PI(18:2(9Z,12Z)/0:0) | C27H49O12P | |
| PI(8:0/9:0) | C9H17O11PC8H15OC9H17O | |
| Picolinic acid | C6H5NO2 | C10164 |
| Pipecolate | C6H11NO2 | C00408 |
| Piperidine | C5H11N | C01746 |
| p-Methoxystilbene | C15H14O | C14679 |
| Prenyl caffeate | C14H16O4 | C10487 |
| Primuletin | C15H10O3 | |
| PS(0:0/13:5) | C6H12NO8PHC13H15O | |
| PS(9:0/14:4) | C6H12NO8PC9H17OC14H19O | |
| PS(9:0/14:5) | C6H12NO8PC9H17OC14H17O | |
| Putrescine | C4H12N2 | C00134 |
| Pyruvate | C3H4O3 | C00022 |
| Quercetin 3'-isobutyrate | C19H16O8 | |
| Quinic acid | C7H12O6 | C00296 |
| Rhamnetin 3'-glucuronide-3,5,4'-trisulfate | C22H20O22S3 | |
| Rhamnose | C6H12O5 | C00507 |
| S2, 3-Epoxysqualene | C30H50O | C01054 |
| S3 Hydroxy-3-methylglutaryl-CoA | C27H44N7O20P3S | C00356 |
| S-Adenosyl-L-methionine | C15H22N6O5S | C00019 |
| Salicin 6-phosphate | C13H19O10P | C06188 |
| Salicylate | C7H6O3 | C00805 |
| Salicylic acid beta-D-glucose ester | C13H16O8 | |
| Scoparone | C11H10O4 | C09311 |

| | | |
|---|---|---|
| Scopoletin | C10H8O4 | C01752 |
| Sebacic acid | C10H18O4 | C08277 |
| Selenohomocystine | C8H16N2O4Se2 | C05707 |
| Se-Methylselenocysteine | C4H9NO2Se | C18905 |
| Serotonin | C10H12N2O | C00780 |
| Shikimate | C7H10O5 | C00493 |
| Shikimate 3-phosphate | C7H11O8P | |
| Sinapate | C11H12O5 | C00482 |
| Sinapoyl aldehyde | C11H12O4 | C05610 |
| Sinapoylagmatine | C16H24N4O4 | |
| Sinapoylhydroxyagmatine | C16H24N4O5 | |
| Sinapyl alcohol | C11H14O4 | C02325 |
| Spermic acid 2 | C10H20N2O4 | |
| Spermidine | C7H19N3 | C00315 |
| Sphinganine 1-phosphate | C18H40NO5P | C01120 |
| Squalene | C30H50 | C00751 |
| Stearic acid | C18H36O2 | C01530 |
| Stearoyl-CoA | C39H70N7O17P3S | C00412 |
| Sterculic acid | C19H34O2 | |
| Stigmasterol | C29H48O | |
| Strigol | C19H22O6 | C09190 |
| Succinate | C4H6O4 | C00042 |
| Succinyl-CoA | C25H40N7O19P3S | C00091 |
| Sugeonyl acetate | C17H24O3 | C17506 |
| Syringic acid | C9H10O5 | C10833 |
| Syringin | C17H24O9 | |
| Trans-2, 3, 4-Trimethoxycinnamate | C12H14O5 | |
| trans-Cinnamate | C9H8O2 | C00423 |
| trans-Cinnamoyl beta-D-glucoside | C15H18O7 | C04164 |
| trans-trans-Farnesyl-diphosphate | C15H28O7P2 | C00448 |
| Traumatic acid | C12H20O4 | C16308 |
| Traumatin | C12H20O3 | C16309 |
| TRIBOA-Gly | C14H17NO10 | C18062 |
| Trigonelline | C7H7NO2 | C01004 |
| TRIMBOA | C9H9NO6 | |
| TRIMBOA-Gly | C15H19NO11 | |
| Tryptamine | C10H12N2 | C00398 |
| Tyramine | C8H11NO | C00483 |
| Uridine diphosphate glucose | C15H24N2O17P2 | |
| Vanillylamine | C8H11NO2 | C16666 |
| Violaxanthin | C40H56O4 | |
| Vitamin K | C31H46O2 | |
| Zeatin | C10H13N5O | C00371 |
| Zeaxanthin | C40H56O2 | C06098 |
| Zeaxanthin diglucoside | C52H76O12 | C15969 |
