## Supplemental Table 2 for "Hemibiotrophic fungal pathogen induces systemic susceptibility and systemic shifts in wheat metabolome and microbiome composition"

| 4 dpi | | |
|---|---|---|
| **Molecular formula** | **Compound name** | **Significant at** |
| C15H14O7 | (+)-Gallocatechin | CS_m-adj ***, CS_m-loc ***, CS_Zt-adj ***, CS_Zt-loc ** |
| C6H12O4 | (R)-Mevalonate | loc_CS-Zt * |
| C12H18O4 | 12-Hydroxy-JA | CS_m-loc * |
| C18H29NO5 | 12-Hydroxy-jasmonic acid-Ile | Ob_m-loc * |
| C5H11O7P | 1-Deoxy-D-xylulose 5-phosphate | loc_Ob-Zt * |
| C17H22O10 | 1-O-Sinapoyl-beta-D-glucose | adj_CS-m **, CS_m-adj **, m_CS-adj * |
| C8H9NO | 2-Phenylacetamide | adj_CS-m *, CS_m-adj * |
| C9H8O6 | 3-(2-Carboxyethenyl)-cis,cis-muconate | adj_CS-Zt ***, Ob_Zt-loc *** |
| C3H7O7P | 3-Phospho-D-glycerate | CS_Zt-loc **, CS_m-loc **, CS_m-adj * |
| C14H12O2 | 4,4'-Dihydroxystilbene | adj_Ob-m *** |
| C36H36O11 | 4'-O-Methylneobavaisoflavone 7-O-(2''-p-coumaroylglucoside) | adj_CS-Zt *** |
| C18H34O6 | 9,10-Dihydroxystearic acid | loc_Ob-m **, loc_CS-m * |
| C15H22O3 | Abscisic alcohol | adj_Ob-Zt * |
| C18H30O2 | alpha-Linolenic acid | Ob_m-adj ** |
| C40H56O3 | Antheraxanthin | adj_CS-Zt **, C_Zt-adj ** |
| C6H8O6 | Ascorbate | loc_Ob-Zt * |
| C21H20O11 | Aureusidin 6-O-glucoside | Ob_Zt-loc * |
| C5H11NO2 | Betaine | Ob_Zt-adj **, Ob_Zt-loc *, Ob_m-loc *, Ob_m-adj * |
| C28H48O6 | Brassinolide | adj_Ob-m * |
| C28H48O | Campesterol | CS_Zt-adj *, adj_Ob-m *, adj_CS-Zt * |
| C16H18O9 | Chlorogenate | CS_Zt-loc ** |
| C9H10O | Cinnamyl alcohol | CS_m-loc ***, CS_Zt-adj ***, CS_Zt-loc ** |
| C14H12O | cis-Stilbene oxide | adj_Ob-Zt * |
| C14H20N4O3 | Coumaroylhydroxyagmatine | loc_CS-Zt *, Ob-Zt-adj * |
| C14H16O7 | de-Hypoxanthine futalosine | loc_CS-m * |
| C8H7NO4 | DIBOA | CS_Zt-adj **, CS_m-loc **, CS_Zt-loc *, CS_m-adj * |
| C20H18O8 | Diferulic acid | loc_CS-m ***, CS_m-loc * |
| C9H9NO5 | DIMBOA | CS_Zt-loc ** |
| C15H19NO10 | DIMBOA-Gly | CS_Zt-adj ** |
| C8H11NO2 | Dopamine | CS_Zt-adj * |
| C8H14O4 | Ethyladipic acid | CS_m-loc **, CS_Zt-adj **, CS_m-adj *, CS_Zt-loc * |
| C15H22N4O3 | Feruloylagmatine | Zt_Ob-adj * |
| C15H22N4O4 | Feruloylhydroxyagmatine | Ob_Zt-adj *, loc_CS-Zt |
| C14H20N2O3 | Feruloylputrescine | Ob_Zt-adj * |
| C4H4O4 | Fumarate | CS_Zt-loc *, loc_CS-Zt *, CS_m-adj *, CS_m-loc * |
| C10H17N3O6S | Glutathione | loc_Ob-Zt * |
| C14H17NO8 | HBOA-Gly | CS_Zt-adj * |

| | | |
|---|---|---|
| C9H9NO4 | HMBOA | CS_Zt-loc *, loc_CS-Zt * |
| C15H19NO9 | HMBOA-Gly | CS_Zt-adj *, CS_m-adj * |
| C43H64N8O17 | Hordatine D + 2 hex | loc_CS-Zt ** |
| C28H38N8O5 | Hydroxy-hordatine A | loc_CS-m *, loc_Ob-m *, m_CS-loc * |
| C8H7NO | Indolin-2-one | Ob_Zt-loc **, Ob_Zt-adj **, Ob_m-loc *, Ob_m-adj * |
| C21H20O10 | Kaempferol 7-rhamnoside | loc_CS-Zt * |
| C27H30O15 | Kaempferol-3-glucoside-3-rhamnoside | loc_Ob-Zt **, loc_CS-Zt *, Ob_Zt-loc * |
| C3H7NO2 | L-Alanine | CS_m-adj * |
| C6H14N4O2 | L-Arginine | CS_Zt-loc * |
| C4H8N2O3 | L-Asparagine | CS_m-adj * |
| C4H7NO4 | L-Aspartate | CS_m-loc * |
| C4H6O5 | L-Malate | CS_m-adj **, CS_Zt-loc * |
| C5H11NO2S | L-Methionine | CS_m-adj * |
| C4H9NO3 | L-Threonine | CS_m-adj * |
| C19H34O5 | Methyl 9,12-dihydroxy-13-oxo-10-octadecenoate | CS_Zt-adj **, CS_m-loc **, CS_Zt-loc *, CS_m-adj * |
| C14H28O2 | Myristic acid | loc_CS-Zt * |
| C26H28O15 | Neocarlinoside | Ob_Zt-loc ***, Ob_m-loc *, Ob_m-adj * |
| C18H36O2 | Octadecanoic acid | loc_Ob-Zt *, loc_Ob-m * |
| C17H20O9 | O-Feruloylquinate | loc_Ob-m * |
| C21H37O7P | PA(18:3(6Z,9Z,12Z)/0:0) | loc_CS-Zt ***, CS_Zt-loc ***, Zt_CS-loc *** |
| C15H18O8 | p-Coumaroyl-D-glucose | CS_m-adj *, CS_m-loc *, CS_Zt-adj * |
| C24H43O9P | PG(18:3(6Z,9Z,12Z)/0:0) | Ob_Zt-adj ***, Ob_m-adj ** |
| C13H10O3 | Phenylsalicylate | CS_m-loc **, Zt_CS-loc ** |
| C9H13NO | Phenylpropanolamine | CS_m-adj **, CS_Zt-loc **, CS_m-loc **, CS_Zt-adj ** |
| C4H12N2 | Putrescine | CS_m-adj * |
| C3H4O3 | Pyruvate | loc_Ob-Zt * |
| C15H22N6O5S | S-Adenosyl-L-methionine | CS_Zt-adj * |
| C7H11O8P | Shikimate 3-phosphate | Ob_Zt-loc ** |
| C7H10O5 | Shikimate | CS_Zt-loc * |
| C16H24N4O4 | Sinapoylagmatine | loc_CS-Zt **, Ob_Zt-adj *, Zt_Ob-adj * |
| C30H50O | Squalene 2,3-epoxide | adj_CS-Zt *, CS_Zt-adj * |
| C30H50 | Squalene | CS_m-loc **, CS_Zt-adj ** |
| C29H48O | Stigmasterol | adj_CS-Zt *, Ob_Zt-adj *, Ob_Zt-loc * |
| C17H24O3 | Sugeonyl acetate | CS_Zt-adj **, CS_m-loc * |
| C10H12N2 | Tryptamine | loc_Ob-m ** |
| C8H11NO | Tyramine | CS_m-adj *, CS_Zt-adj *, CS_Zt-loc * |
| C15H24N2O17P2 | Uridine diphosphate glucose | CS_Zt-loc * |
| C40H56O4 | Violaxanthin | loc_CS-Zt * |

| 8 dpi | | |
|---|---|---|
| Molecular formula | Compound name | Significant at |
| C15H14O7 | (+)-Gallocatechin | CS_m-adj ***, CS_m-loc ***, CS_Zt-loc **, CS_Zt-adj ** |
| C18H34O2 | (9Z)-Octadecenoic acid | adj_Ob-m **, adj_CS-Zt *, adj_CS-m * |
| C8H9NO | 2-Phenylacetamide | CS_m-loc *** |
| C30H28O12 | 4,2',3',4'-Tetrahydroxychalcone 4'-O-(2”-O-p-coumaroyl) glucoside | Ob_Zt-adj ** |
| C14H12O2 | 4,4'-Dihydroxystilbene | loc_CS-Zt * |
| C8H8O3 | Methylsalicylate | loc_Ob-Zt * |
| C26H32O12 | 5-Hydroxy-6,7,3',4',5'-pentamethoxyflavanone 5-O-rhamnoside | adj_CS-m **, adj_Ob-m ** |
| C18H34O6 | 9,10-Dihydroxystearic acid | loc_CS-m **, m_CS-loc ** |
| C20H30O | 9-cis-Retinol | Ob-Zt-loc * |
| C9H16O3 | 9-hydroxy-5Z-nonenoic acid | CS_Zt-loc *, CS_Zt-adj *, adj_Ob-m * |
| C29H50O2 | alpha-Tocopherol | adj_Ob-m *, adj_CS-m * |
| C40H56O3 | Antheraxanthin | adj_CS-m **, adj_Ob-m *, adj_CS-Zt *, Zt_CS-loc * |
| C5H11NO2 | Betaine | Ob_m-loc * |
| C9H10O3 | Caffeyl alcohol | CS_Zt-adj * |
| C28H48O | Campesterol | adj_Ob-m **, adj_CS-m **, adj_Ob-Zt *, adj_CS-Zt * |
| C16H18O9 | Chlorogenate | CS_m-adj **, loc_Ob-Zt ** |
| C10H16O2 | Chrysanthemic acid | CS_Zt-loc * |
| C9H10O | Cinnamyl alcohol | CS_Zt-adj ***, CS_m-adj ***, CS_m-loc ***, CS_Zt-loc ***, adj_CS-m * |
| C6H6O6 | cis-Aconitate | Ob_Zt-adj *, loc_CS-Zt*, loc_CS-m * |
| C6H8O7 | Citrate | loc_CS-Zt **, loc_Ob-m ** |
| C12H14O4 | Coniferyl acetate | CS_m-loc **, CS_Zt-loc **, adj_Ob-m ** |
| C14H20N4O3 | Coumaroylhydroxyagmatine | loc_CS-Zt *, Zt_Ob-loc * |
| C14H16O7 | de-Hypoxanthine futalosine | loc_Ob-Zt **, loc_Ob-Zt * |
| C6H13O9P | D-Glucose 1-phosphate | loc_CS-Zt **, loc_Ob-Zt * |
| C8H7NO4 | DIBOA | CS_Zt-adj **, Ob_Zt-loc * |
| C14H14O3 | Dihydroresveratrol | CS_m-loc * |
| C9H9NO5 | DIMBOA | loc_Ob-Zt * |
| C8H11NO2 | Dopamine | CS_m-loc ** |
| C8H14O4 | Ethyladipic acid | CS_m-loc ***, CS_Zt-adj *, CS_Zt-loc *, CS_m-adj * |
| C15H28O7P2 | Farnesyl diphosphate | loc_CS-Zt **, CS_Zt-loc * |
| C15H22N4O4 | Feruloylhydroxyagmatine | loc_CS-Zt ** |
| C14H20N2O3 | Feruloylputrescine | loc_CS-Zt * |
| C15H10O3 | Flavonol | CS_Zt-loc **, CS_m-loc **, adj_Ob-m ** |
| C19H24O6 | Gibberellin | loc_Ob-Zt *, CS_m-adj * |
| C8H7NO3 | HBOA | Ob_m-loc * |
| C14H17NO8 | HBOA-Gly | CS_Zt-loc * |
| C10H11NO5 | HDMBOA | loc_Ob-Zt ** |

| | | |
|---|---|---|
| C9H9NO4 | HMBOA | CS_Zt-loc **, CS_m-loc **, CS_m-adj **, loc_CS-m **, CS_Zt-adj ** |
| C15H19NO9 | HMBOA-Gly | loc_Ob-Zt **, CS_m-adj *, CS_Zt-adj *, loc_Ob-Zt *, CS_m-loc* |
| C8H7NO | Indolin-2-one | Ob_m-loc * |
| C27H30O14 | Kaempferol 3-rhamnoside-(1->2)-rhamnoside | loc_CS-Zt *, loc_Ob-Zt *, CS_m-loc * |
| C27H30O15 | Kaempferol-3-glucoside-3-rhamnoside | loc_CS-Zt **, loc_CS-m * |
| C4H8N2O3 | L-Asparagine | CS_Zt-adj **, CS_m-adj * |
| C4H7NO4 | L-Aspartate | CS_m-adj * |
| C6H10O6 | L-Galactono-1,4-lactone | loc_Ob-Zt * |
| C5H9NO4 | L-Glutamate | CS_m-loc **, CS_Zt-adj * |
| C9H11NO2 | L-Phenylalanine | adj_Ob-m *** |
| C4H9NO3 | L-Threonine | CS_Zt-adj **, CS_Zt-loc *, CS_m-adj *, adj_CS-Zt * |
| C9H11NO3 | L-Tyrosine | CS_Zt-adj * |
| C13H18N2O3 | N-Caffeoylputrescine | loc_CS-Zt ** |
| C19H18N2O3 | N-Coumaroyl serotonin | Ob_m-loc * |
| C26H28O15 | Neocarlinoside | Ob_Zt-adj *, Ob_m-loc *, Ob_m-adj * |
| C4H4O5 | Oxaloacetate | loc_CS-m ** |
| C15H18O8 | p-Coumaroyl-D-glucose | CS_Zt-adj *, CS_m-loc * |
| C9H13NO | Phenylpropanolamine | CS_Zt-adj **, CS_Zt-loc **, adj_CS-m **, CS_m-adj **, Zt_CS-loc ** |
| C3H5O6P | Phosphoenolpyruvate | Ob_Zt-adj * |
| C5H9N | Piperideine | CS_Zt-loc * |
| C4H12N2 | Putrescine | loc_CS-Zt * |
| C7H12O6 | Quinic acid | adj_Ob-Zt *, CS_Zt-adj * |
| C15H22N6O5S | S-Adenosyl-L-methionine | CS_Zt-loc *, CS_Zt-adj *, CS_m-adj *, CS_m-loc * |
| C13H16O8 | Salicylic acid-D-glucose ester | Ob_m-loc * |
| C10H8O4 | Scopoletin | adj_CS-m * |
| C10H18O4 | Sebacic acid | CS_Zt-loc *, Zt_CS-loc * |
| C16H24N4O4 | Sinapoylagmatine | loc_CS-Zt * |
| C11H14O4 | Sinapyl alcohol | CS_Zt-adj * |
| C10H20N2O4 | Spermic acid 2 | loc_Ob-Zt * |
| C18H40NO5P | Sphinganine 1-phosphate | adj_Ob-m **, adj_CS-m **, adj_CS-Zt *, CS_m-adj *, CS_m-loc * |
| C30H50O | Squalene 2,3-epoxide | adj_Ob-m *, adj_CS-Zt *, adj_CS-m * |
| C30H50 | Squalene | adj_CS-Zt ***, CS_Zt-adj ** |
| C19H34O2 | Sterculic acid | Ob_Zt-loc **, Ob_m-loc *, adj_CS-Zt *, adj_Ob-m *, adj_Ob-Zt *, adj_CS-m * |
| C17H24O3 | Sugeonyl acetate | CS_Zt-adj ***, loc_CS-Zt ***, loc_Ob-Zt *, CS_m-adj *, CS_Zt-loc * |
| C9H10O5 | Syringic acid | Zt_Ob-adj **, CS_m-adj **, loc_CS-Zt * |
| C17H24O9 | Syringin | loc_Ob-Zt **, loc_CS-m **, loc_CS-Zt **, loc_Ob-Zt ** |
| C15H18O7 | trans-Cinnamoyl beta-D-glucoside | loc_Ob-Zt * |
| C31H46O2 | Vitamin K | adj_Ob-m * |
